# Benchmarking algorithms for joint integration of unpaired and paired single-cell RNA-seq and ATAC-seq data

**DOI:** 10.1101/2023.02.01.526609

**Authors:** Michelle Y. Y. Lee, Klaus H. Kaestner, Mingyao Li

## Abstract

Single-cell RNA-sequencing (scRNA-seq) measures gene expression in single cells, while single-nucleus ATAC-sequencing (snATAC-seq) enables the quantification of chromatin accessibility in single nuclei. These two data types provide complementary information for deciphering cell types/states. However, when analyzed individually, scRNA-seq and snATAC-seq data often produce conflicting results regarding cell type/state assignment. In addition, there is a loss of power as the two modalities reflect the same underlying cell types/states. Recently, it has become possible to measure both gene expression and chromatin accessibility from the same nucleus. Such paired data make it possible to directly model the relationships between the two modalities. However, given the availability of the vast amount of single-modality data, it is desirable to integrate the paired and unpaired single-modality data to gain a comprehensive view of the cellular complexity. Here, we benchmarked the performance of seven existing single-cell multi-omic data integration methods. Specifically, we evaluated whether these methods are able to uncover peak-gene associations from single-modality data, and to what extent the multiome data can provide additional guidance for the analysis of the existing single-modality data. Our results indicate that multiome data are helpful for annotating single-modality data, but the number of cells in the multiome data is critical to ensure a good cell type annotation. Additionally, when generating a multiome dataset, the number of cells is more important than sequencing depth for cell type annotation. Lastly, Seurat v4 is the best at integrating scRNA-seq, snATAC-seq, and multiome data even in the presence of complex batch effects.

## Background

Over the past ten years, hundreds of single-cell RNA-seq (scRNA-seq) (for transcript abundance in single cells) or single-nucleus ATAC-seq (snATAC-seq) (for chromatin accessibility in single nuclei) have been produced by laboratories worldwide, leading to the discovery of new cell types and regulatory circuits. In addition, by applying single-cell assays to two-state models such as the comparison between control and mutant tissues, changes in gene expression or chromatin accessibility caused by a gene mutation could be analyzed at the cell type-specific level easily for the first time. Unfortunately, each single-modality dataset measures either the gene expression or the chromatin accessibility of a given cell. Although the two datasets are generated from the same cell population, they measure different cells. Most of the time, the two experimental modalities result in the identification of similar cell types, as the promoters of highly expressed genes used to define cell types at the transcript levels are frequently also identified as highly accessible by the ATAC-seq modality. However, there are situations in which the two profiles are discordant. In these situations, simultaneous, joint profiling of gene expression and chromatin accessibility is paramount for resolving inconsistency and revealing novel cell types and states that show modality-specific features. Moreover, the joint profiling of gene expression and chromatin accessibility of the same exact cells offers the most direct link between *cis*-regulatory elements and their target genes [1].

Recently, the simultaneous determination of both transcript levels and chromatin state in the same nucleus has become possible, using so-called “multi-omics” approaches. An example is the 10x Genomics single cell Multiome ATAC + gene expression technology [2]. Multi-omics datasets are clearly superior at refining cell types and revealing gene regulatory networks [1]. However, it is not practical to repeat all prior studies of interest performed using the single-modality assays with the multiome approaches, as frequently precious samples are either no longer available or funding is limited. Therefore, it is highly desirable to integrate pre-existing single-modality scRNA-seq and snATAC-seq datasets with multiome data generated subsequently using the newer technology to achieve more accurate cell type annotations.

Several methodologies have been developed for multi-omic data integration. Here, we refer to multi-omic integration as the integration of RNA-seq and ATAC-seq profiles measured in single cells, either with or without the guidance of multiome data. These methods attempt to align cells profiled by separate technologies and project them into one common low-dimensional space to ensure consistent cell type calling. However, we still lack an objective evaluation of whether the addition of the multiome data improves the annotation of single-modality datasets. Furthermore, some of the methods try to impute the missing modality for the single-modality datasets and identify peak-gene pairs using these ‘pseudo-paired’ datasets. Thus, it is still uncertain if the imputed missing modality can truly provide additional biological insights to the same degree as provided by the experimentally produced multiome datasets. Finally, given the availability of many methods for multi-omic data integration, at present, we do not know which method performs the best when integrating all three data types.

The current multi-omic integration methods can be divided into two categories. Methods in the first category perform multi-omic integration using only the single-modality datasets, aiming to find a mapping between gene expression profiles and chromatin accessibility states to create an aligned space that explains both modalities; we call these approaches ‘unpaired integration’. Representative methods in this category include Seurat version 3 (Seurat v3) [3], which performs canonical correlation analysis (CCA) to align experimentally measured gene expression with pseudo-gene expression obtained from chromatin accessibility. One example of pseudo-gene expression is the gene activity score, calculated by summing up peak counts within the gene body plus 2kb upstream in the ATAC-seq data. LIGER [4] also uses the gene expression and gene activity score to obtain shared features between the two modalities and then derives a low-dimensional embedding through a non-negative matrix factorization approach. FigR [5] aligns the snATAC-seq and scRNA-seq data using a CCA-based approach.

In addition, it provides matching of snATAC-seq and scRNA-seq cells, which enables the identification of *cis*-regulatory elements similar to paired multiome data. BindSC [6] goes beyond the simple construction of gene activity scores. Instead, bindSC uses a bi-directional CCA to empirically construct a cell-by-gene matrix for the snATAC-seq cells that preserve its similarity with the ATAC-seq input and simultaneously maximizes the correlation with the scRNA-seq matrix it is being integrated with.

Methods in the second category encompass more recent approaches that incorporate information from multiome cells and integrate all three data types for a more comprehensive exploration of cellular identities; we term these approaches ‘multiome-guided integration’. Representative methods in this category include Seurat version 4 (Seurat v4) [7], an approach that first learns a low-dimensional representation of the cells profiled by the multiome methodology using both the RNA-seq and ATAC-seq profiles by weighted nearest neighbors (WNN) analysis [7]. Subsequently, the two single-modality datasets are projected onto the WNN embedding space in a supervised manner. MultiVI [8] and Cobolt [9] use a deep-learning approach called ‘variational autoencoder’ to embed all three data types. Both methods employ the encoder-decoder system to learn a low-dimensional representation of the data. Specifically, two encoders and two decoders are set up, one for each modality. However, there are different model choices. MultiVI assumes a negative binomial distribution for the RNA-seq data and a Bernoulli distribution for the ATAC-seq data, while Cobolt assumes a Multivariate Normal distribution for both modalities. Furthermore, the two methods integrate the modality-specific representation for the paired cells differently. MultiVI first aligns the two embeddings through a symmetric Kullback-Leibler (KL) divergence loss and then obtains an average of the two embeddings. On the other hand, Cobolt simply multiplies the two embeddings to represent the paired cells, while the representation of the unpaired cells is first generated by the corresponding encoder and refined using a linear transformation to ensure enough similarity between the RNA-seq derived embedding and the ATAC-seq derived embedding.

All methods described above aim to project cells from different data types into one shared space to facilitate the identification of cell types through clustering. Nevertheless, a common goal for studies profiling chromatin accessibility and gene expression at the single-cell level is to understand cell type-specific *cis*-regulatory logic. Since the two single-modality datasets are generated from different cells in a given population, albeit representing the same cell types, the single-modality datasets cannot be naïvely combined to test for association between chromatin accessibility and gene expressions. Therefore, multiple efforts have attempted to impute the missing modality for the single-modality datasets, aiming to computationally generate paired profiles similar to those measured experimentally by the multiome technology. Some methods mentioned above, e.g., Seurat v3, FigR, bindSC, Seurat v4, and MultiVI, are capable of this task. However, an objective evaluation of how reliable the *in-silico* imputed profiles are compared to what is directly measured by the paired multiome technologies is still lacking. Therefore, we aimed to conduct an extensive benchmarking analysis to evaluate the above-mentioned methods by addressing two important questions. First, do multiome data help the integration of single-modality datasets? Second, what is the best computational method for the integration of scRNA-seq, snATAC-seq, and multiome data?

## Results

### Overview of the benchmarking scheme and evaluation strategies

The overall workflow of our benchmarking evaluations is summarized in Figure 1. Figure 1A illustrates our approach to evaluate whether multiome data integration can improve the value of single-modality datasets, while Figure 1B outlines how we assess the effectiveness of each integration methods, at various conditions of the multiome dataset. To answer the proposed questions, we simulated situations where all three data types are available by using two publicly available multiome datasets [10, 11]. The first multiome dataset [10] profiled 10,085 peripheral blood mononuclear cells (PMBCs) and represents a simple biological system, because PBMCs can be easily divided into seven well-separated cell types (Supplementary Figure 1A). The second dataset profiled bone marrow mononuclear cells (BMMC) [11], an example of highly complex cell populations. BMMCs are closely related to each other transcriptionally, and contain, for example, myeloid progenitors and their closely related descendants, CD16+ and CD14+ monocytes (Supplementary Figure 1B). The individual BMMC cell types are therefore much harder to separate compared to the PBMC populations, thus allowing us to thoroughly evaluate the performance of each method in both simple and complex biological systems. Moreover, the BMMC dataset is composed of samples generated from four research sites and nine donors [12], which enables the analysis of batch effects and technical replicates.

**Figure 1:**
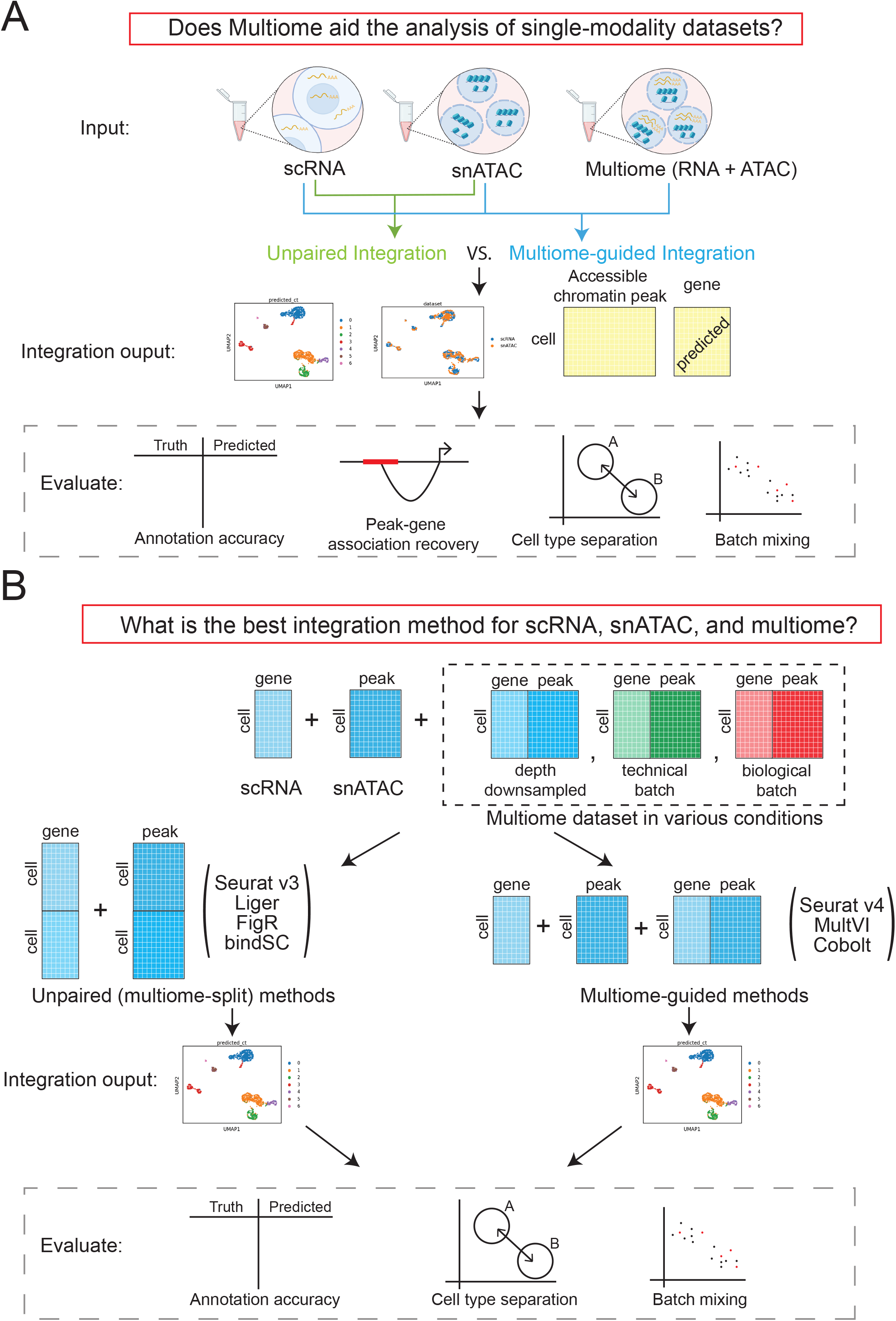
Outline of the benchmarking evaluations. (A) Scheme to evaluate if multiome data help the integration of single-modality data. (B) Scenarios simulated to evaluate multi-omic integration methods.

We evaluated four popular unpaired integration methods (Seurat v4, LIGER, FigR, bindSC), and three multiome-guided integration methods (Seurat v4, MultiVI, and Cobolt). To account for the increased power resulting from the larger absolute number of cells employed during the integration process by the multiome-guided methods, we created another scenario termed ‘unpaired (multiome-split)’ in which the RNA-seq and ATAC-seq data from the multiome samples were treated as independent datasets and appended to the single-modality datasets. This category again includes the four unpaired-integration methods, the only difference being that the single-modality datasets now include additional single-modality cells that were converted from the multiome cells.

To evaluate the performance of each method for cell type identification, we performed Louvain clustering [13] on the integrated embedding. For methods capable of missing modality imputation, we imputed gene expression using snATAC-seq profiles. We then evaluated the integration results in four aspects as shown in Figure 1A. Specifically, we evaluated cell type annotation accuracy using two metrics: Adjusted Rand Index (ARI) [14] and Normalized Mutual Information (NMI) [15]. Both metrics range from 0 to 1, with 1 being the best. The detailed approach is described in the Methods section. The accuracy of cell type annotation depends on the number of cell clusters identified; therefore, an additional way to measure data integration quality is via the accuracy of cell type separation. Using the ground-truth annotation, we evaluated how well cells of different identities are separated, using a cell type specific average silhouette width (ASW) [16] and a cell type Local Inverse Simpson’s Index (cLISI) [17, 18]. Furthermore, because the three data types could have technology-specific differences, we used a batch ASW [16] and the k-nearest neighbor batch effect test (kBET) [16] to measure batch-mixing of the integrated results. These four measurements were normalized to be in the range of 0 and 1 in which 1 is the best result, namely high separation between cell types and complete mixing of data batches.

We also evaluated the quality of ‘peak to gene pair’ predictions by assessing the accuracy of assigning an ATAC-seq peak to a specific gene. Using the measured ATAC-seq and imputed RNA-seq data, we computed the percentage of significant peak-gene pairs recovered as compared to a ground truth obtained using all cells in the multiome dataset. To penalize for the presence of false positives reported by the data integration methods, we also calculated an F1 score [15], which normalized the absolute percent recovery of the true peak-gene pairs by the occurrence of false positive and false negative relationships.

### Do Multiome data improve the annotation of single-modality datasets?

#### PBMC

To answer if multiome data improve the analysis of single-modality datasets (scRNA-seq and scATAC-seq), we first simulated the situation with 1,000 scRNA-seq cells and 1,000 snATAC-seq cells based on the PBMC data. These single-modality cells were integrated using each of the four unpaired integration methods. To evaluate if multiome data improve the analysis of single-modality datasets, we considered the situation where we have a multiome dataset, potentially with different numbers of cells (e.g., 1000, 3000, or 8000). These multiome data were integrated with the single-modality datasets using the multiome-guided methods. However, because the number of cells used during clustering and gene expression imputation impacts the clustering accuracy and peak-gene association identification, we ran the unpaired integration methods again, this time treating the multiome dataset as single-modality cells and adding them to the existing single-modality data. Here, any increase in performance is solely caused by the increase in cell number; the results from these evaluations are labeled as the ‘unpaired (multiome-split)’ category. For each simulation, we randomly drew the cells from the 10,085 PBMC dataset and each condition was repeated five times. The parameters used for this simulation are summarized in Figure 2A.

**Figure 2:**
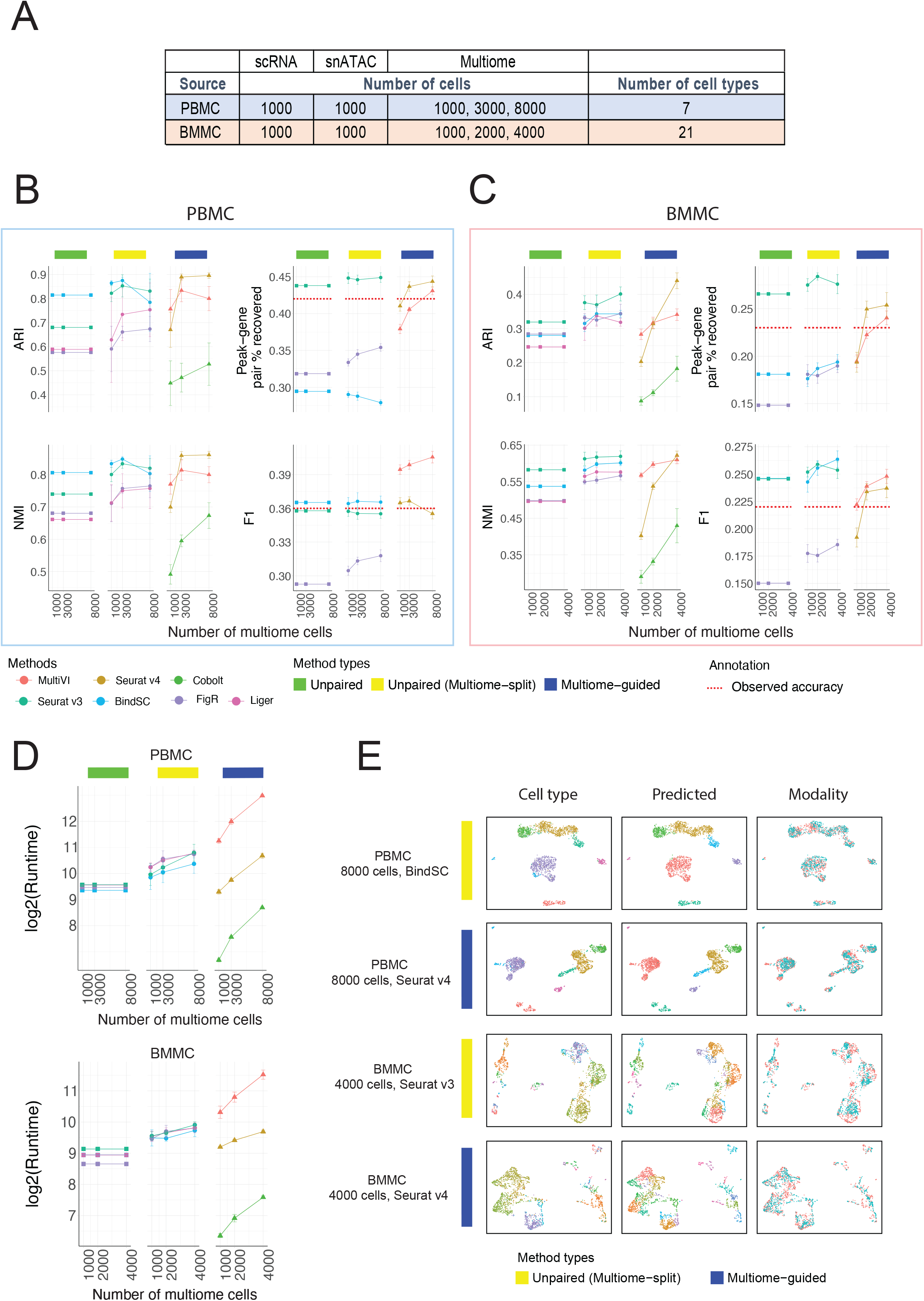
Comparison of integration performance without vs. with multiome cells. (A) The number of cells and cell types for each simulated dataset using the PBMC or BMMC multiome data as the ground truth. (B – C) Performance of cell type annotation and peak-gene association recovery in the PBMC-based simulations (B) and BMMC-based simulations (C). ARI and NMI measure agreement between predicted cell type and ground-truth labels. Peak-gene pair % recovered is the percentage of peak-gene pairs correctly identified comparing to the ground-truth list calculated using 10,412 paired PBMC cells (B) and 6,740 BMMC cells (C). F1 is the prediction accuracy normalized by the number of false positives and false negatives. Dashed line shows the percent recovery and F1 score calculated using 1,000 multiome cells. Error bar is mean ± standard deviation. (D) Runtime measured in seconds, for each method, in log2 scale. Error bar is mean ± standard deviation. (E) UMAP projection using integrated embedding for a select number of methods. UMAP projection for the other methods are shown in Supplementary Figures 3 (PBMC) and 4 (BMMC).

The evaluation result for each method is summarized in Figure 2B. Without the incorporation of multiome data, the cell type annotation accuracy was already good, being 0.81 in ARI and 0.81 in NMI when integrating the unpaired data using the bindSC program (Figure 2B). Surprisingly, in the presence of 1,000 multiome cells, the multiome-guided approaches performed worse than simply integrating the data from the 2,000 single-modality cells by themselves (Figure 2B). This unexpected result is likely caused by the fact that 1,000 multiome cells alone do not achieve good cell type separation, which is a critical requirement for the multiome-guided methods to succeed. However, when we used 3,000 or 8,000 multiome cells, Seurat v4, one of the multiome-guided methods, achieved the best results in terms of cell type annotation (Figure 2B). Furthermore, when comparing the multiome-guided results with unpaired (multiome-split) results, the performance of Seurat v4 remained higher when there are 3,000 or 8,000 cells (Figure 2B). Thus, our findings indicate that the multiome data can improve cell type annotation of the single modality datasets, provided that there is a sufficient number of multiome cells available.

Next, we evaluated the performance of each method in predicting peak-gene pairs. Peak-gene pairs are calculated using 1,000 measured chromatin accessibility profiles and the corresponding 1,000 imputed gene expression profiles. Here, we compared predicted peak-gene pairs to the ground-truth list calculated using multiome cells in the full PBMC data. Seurat v3 performed very well at recovering the absolute number of peak-gene pairs, and the incorporation of data from multiome cells through splitting only marginally increased the performance (Figure 2B). BindSC had a slightly better F1 score than Seurat v3, meaning that the Seurat v3 results contained more false positives (Figure 2B). For the multiome-guided methods, the more multiome cells available during gene expression imputation resulted in higher peak-gene pair recovery (Figure 2B). Nevertheless, the incorporation of data from multiome cells using the multiome-guided methods did not perform better than the unpaired methods, with the exception that the F1 score was higher in MultiVI (Figure 2B).

The number of cells used for predicting peak-gene pairs influences the accuracy. To give a general idea of how well the predicted gene expression profiles are, we compared the peak-gene pair identification result to the one obtained using the real paired profiles. We included a red dashed line in Figure 2B to indicate the percentage of peak-gene pair recovery and F1 score calculated using the measured, paired gene expression and chromatin accessibility profiles of the 1,000 cells being evaluated, instead of the gene expression profile imputed from chromatin accessibility. What’s surprising is that the in-silico prediction profile from Seurat v3 revealed a higher percentage of recovered peak-gene pairs and a better F1 score than the measured paired gene expression and chromatin accessibility profile from 1,000 cells. This is likely due to the dropout issue common to single-cell assays and the predicted RNA profile can borrow information from similar cells, thus recovering the trend better. However, we also note that the predicted profiles only recovered less than 45% of the ground-truth list calculated using the full PBMC data with 10,412 cells. Although the predicted profiles are better than the measured gene expression profiles, it is only recovering a small percentage of peak-gene pairs revealed by the experimentally generated multiome dataset.

#### BMMC

Having evaluated the various data integration platforms with the PBMC data, which represent a low-complexity situation with clearly defined major cell types, we next sought to determine how the different methodologies perform when analyzing data from highly complex cell populations, as is the case for bone marrow mononuclear cells (BMMC). Here, to avoid complexity caused by batch differences, we only used 6,740 multiome cells from one sample (site 1 donor 2). We again started with 1,000 scRNA-seq and 1,000 snATAC-seq cells, and then tested the result when incorporating 1000, 2000, and 4000 multiome cells, composed of 21 cell types (Figure 2A). In this biological system, we found that including a larger number of multiome cells improved cell type annotation, with Seurat v3 performing the best among the unpaired (multiome-split) methods (Figure 2C). Among the multiome-guided methods, Seurat v4 achieved the best performance when the input data included 4,000 multiome cells. Remarkably, when we employed data from only 1,000 or 2,000 multiome cells, all multiome-guided methods performed worse than when inputting the multiome data as two separate, unpaired modalities (Figure 2C). A similar trend was observed in the peak-gene pair prediction (Figure 2C). The likely reason causing the poor performance of the multiome-guided methods is the limited quality of multiome data and the high complexity of the biological system being profiled. Note that peak-gene prediction recovery and F1 score obtained via the unpaired Seurat v3 algorithm are still higher than the association calculated from the observed multiome profile indicated by the red dash line in Figure 2C.

#### Comparison of run time and visualization of integration

Another important issue to consider when comparing various computational approaches is the computation time needed to complete a given task. All methods were run with 8 CPU cores and 64GB of RAM. Figure 2D shows the runtime, measured in seconds. Unpaired methods all have similar runtimes, and the increase in the unpaired (multiome-split) category was due to the incorporation of the additional data from multiome experiments. Importantly, the multiome-guided methods vary greatly in runtime and thus costs. Cobolt was the fastest method, but unfortunately, it exhibited comparatively low clustering accuracy and peak-gene recovery. Seurat v4 had a shorter runtime than the unpaired (multiome-split) methods, while MultiVI took the longest to complete the assigned tasks, due to its use of variational autoencoder.

To visually examine the integration results, we generated UMAP plots using the integrated latent embedding and colored the cells by the ground-truth annotation, the predicted identity, and the dataset origin (Figure 2E). We showed the best-performing results from both the unpaired (multiome-split) and multiome-guided categories for each of the PBMC and BMMC simulations. Additional evaluation on cell type separation and batch-mixing are shown in Supplementary Figure 2. Most metrics show method-specific values, meaning the rankings of methods do not change across different numbers of multiome cells. Among the unpaired methods, Seurat v3 is the best at separating cell types in the integrated space, but it has the worst batch mixing result. On the other hand, FigR shows the opposite trend; it ranked the highest for batch mixing, but the lowest for cell type separation. Among the multiome-guided integration methods, MultiVI mixes the batches better while Seurat v4 often results in a higher cell type silhouette score, especially when there is a greater number of multiome cells. We also evaluated the integration results visually, through examining UMAP projection of the integration results as shown in Supplementary Figure 3 for the PBMC simulations and Supplementary Figure 4 for the BMMC simulations. Visually, we do not see drastic differences between methods and there are no methods showing particularly poor cell type separation or batch mixing result. Therefore, we conclude that the incorporation of multiome cells improves cell type annotation when there are enough cells to resolve the cell type heterogeneity in the multiome dataset alone.

### How to spend your sequencing dollars: more cells or increased sequencing depth?

Experimentalists are commonly constrained by budget limitations and need to consider whether sequencing a larger number of cells at low depth or a smaller number of cells at high depth is the more productive approach. To answer this question, we evaluated how the sequencing depth of the multiome approach influences the integration result. Since we know that including multiome data improves cell type annotation for the single-modality datasets, for this analysis, we aimed to evaluate the cell type annotation accuracy of the three data types together. Table 1 shows the sequencing depth of the original multiome samples. To simulate data with lower depths, we down-sampled the reads for both RNA and ATAC profiles to 25%, 50%, 75% of the original data (Figure 3A) and compared these results to the original samples. We performed this experiment on both the PBMC dataset (Figure 3B) and the BMMC dataset (Figure 3C). For the PBMC study, the increase in sequencing depths resulted in an increase in cell type annotation accuracy for all methods, with Seurat v4 achieving the highest ARI and NMI among all methods for 75% and 100% depth. In contrast, when we used the BMMC data set as the input, we noted that when including only 2,000 multiome cells, regardless of sequencing depth, the unpaired method (Seurat v3) performed the best. However, when we included 4,000 cells in the BMMC multiome sample, 50% of read depth was sufficient for Seurat v4 to annotate the cell types most accurately. These conflicting results prompted us to ask whether sequencing depth is less important than cell number.

**Figure 3:**
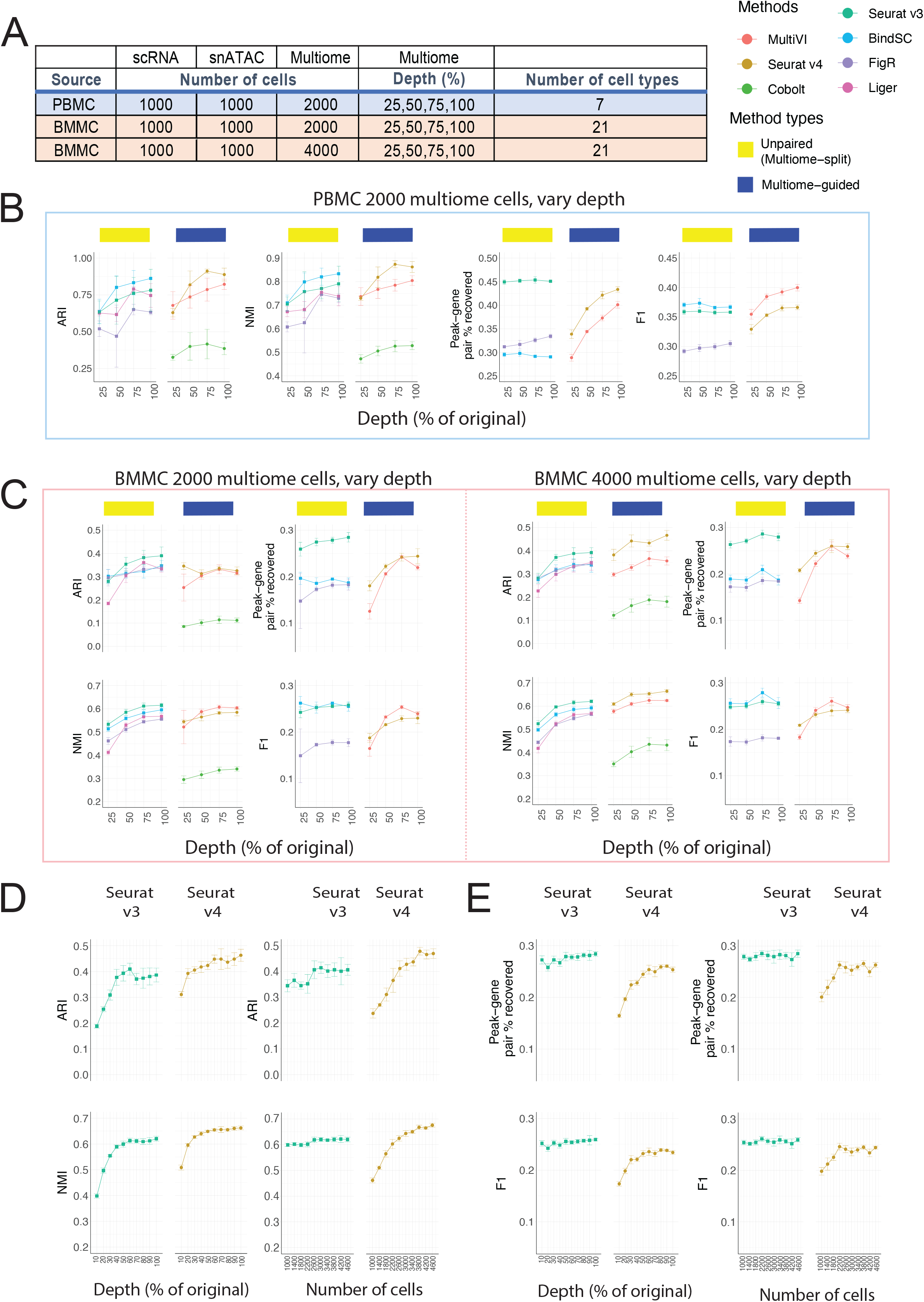
Evaluation of integration performance at varying sequencing depth for multiome cells. (A) Details of the simulation scheme. (B – C) Performance of cell type annotation and peak-gene association recovery in the PBMC-based simulations (B) and BMMC-based simulations (C: left panel, 2,000 multiome cells; right panel, 4,000 multiome cells). ARI and NMI measures agreement between predicted cell type and ground-truth labels. Peak-gene pair % recovered is the percentage of peak-gene pairs correctly identified comparing to the ground-truth list calculated using 10,412 paired PBMC cells (B) and 6,740 BMMC cells (C). F1 is the prediction accuracy normalized by the number of false positives and false negatives. (D) Performance of cell type annotation using Seurat v3 or Seurat v4 at increasing depth or increasing number of cells. (E) Performance of peak-gene association recovery using Seurat v3 or Seurat v4 at increasing depth or increasing number of cells. For all subplots, error bar is mean ± standard deviation.

**Table 1:** Summary of the data used for simulation. Columns are number of cells (n_cells), number of unique genes expressed per cell on average in the RNA profile (nGene_RNA), total counts expressed per cell on average in RNA profile (nCount_RNA), number of unique fragments per cell on average in the ATAC profile (nFrag_ATAC), number of peak counts per cell on average in the ATAC profile (nPeakCount_ATAC).

| Source | n_cells | nGene_RNA | nCount_RNA | nFrag_ATAC | nPeakCount_ATAC |
| --- | --- | --- | --- | --- | --- |
| PBMC | 10085 | 2013 | 4463 | 15510 | 11305 |
| BMMC site 1<br>donor 2 (S1D2) | 6740 | 1365 | 2525 | 11064 | 7512 |
| BMMC site 1<br>or donor 1 | 29486 | 1205 | 2227 | 11798 | 7615 |

To answer this question, we designed another simulation. Given a fixed cost for 1,000,000 RNA-seq reads and 4,000,000 ATAC-seq reads, we used either 400 cells with 100% of the depth (see Table 1), or 10% of the reads for 4,000 cells. Next, we analyzed the datasets using Seurat v3 and Seurat v4, the best-performing method in each category based on Figure 3C. For cell type annotation accuracy, the sequencing depth curve plateaued sooner than the number of cells curve. For Seurat v4, the ARI and NMI did not increase much beyond 60% sequencing depth, while both scores increased consistently as the number of cells increases. Comparing Seurat v3 with Seurat v4, we noted that Seurat v4 performed better when there was 30% sequencing depth given 4,000 cells or 2,600 cells given 100% depth. Therefore, for the accuracy of cell type annotation for integrated data, having more cells is more important than having a higher sequencing depth. Importantly, once a sufficient number of cells has been profiled to capture the complexity of a given sample, the multiome-guided methods, specifically Seurat v4, are the best. Our analysis also demonstrated that the ‘sufficient’ number of cells depends on the complexity of the biological system in question. For PBMC, we see that if the goal is to detect seven distinct cell types, 2,000 cells is already enough. However, for BMMC with its more complex cell type composition at least 2,600 cells are needed.

In addition to the cell type annotation accuracy, we also evaluated recovery of peak-gene association for the 1,000 single-modality ATAC-seq cells when incorporating mulitome samples generated at ten different depths and numbers of cells. We see that Seurat v3 is consistently better than Seurat v4 (Figure 3D). Moreover, the number of cells and sequencing depth did not affect the percentage of peak-gene pair recovery nor the F1 score. This is likely because Seurat v3 predicts RNA expression using a nearest neighbor approach on the integrated space, and the software had enough cells in the scRNA-seq dataset for the prediction, thus changes in the multiome data did not affect the result.

Next, we evaluated cell type separation and batch mixing results as summarized in Supplementary Figure 5. Most metrics increased slightly as sequencing depth increased, but the ranking of methods is similar as described before. Overall, Seurat v4 shows the best separation of cell types in the integrated space, but the mixing of batches is the worst, across sequencing depths. A UMAP projection of each method under each simulated scenario is shown in Supplementary Figures 6-8 for visual comparison.

Overall, we conclude that the number of cells in the multiome data is more critical than sequencing depth for annotating cell types in the integrated data. On the other hand, treating multiome data as unpaired single-modality datasets recovers peak-gene pairs at a higher accuracy.

### Which method is the best at removing batch effects?

It is common that scRNA-seq and snATAC-seq data are generated by different labs or from different individuals than the multiome data. Therefore, another key characteristic for integration methods is whether they can integrate samples displaying batch effects. To answer this question, we leveraged the complex batch structure present in the BMMC dataset. Figure 4A shows the technical batch or biological batch structure we aimed to evaluate, with the multiome cells coming from a different research site, or a different donor. Figure 4B shows the results of cell type annotation accuracy for unpaired integration methods and the multiome-guided methods. We again saw increasing cell type annotation accuracy as the number of multiome cells increased. With 3,000 or more multiome cells, Seurat v4 again was the best-performing method. Seurat v4 is a supervised approach, meaning that the multiome sample serves as a reference to which the single-modality datasets are mapped to. Figure 4B shows that although the multiome sample has strong batch effects (Supplementary Figure 9), the supervised mapping approach resulted in the most accurate cell type annotation. Additional integration results are shown in Supplementary Figure 10 and the UMAP projections are shown in Supplementary Figures 11-12.

**Figure 4:**
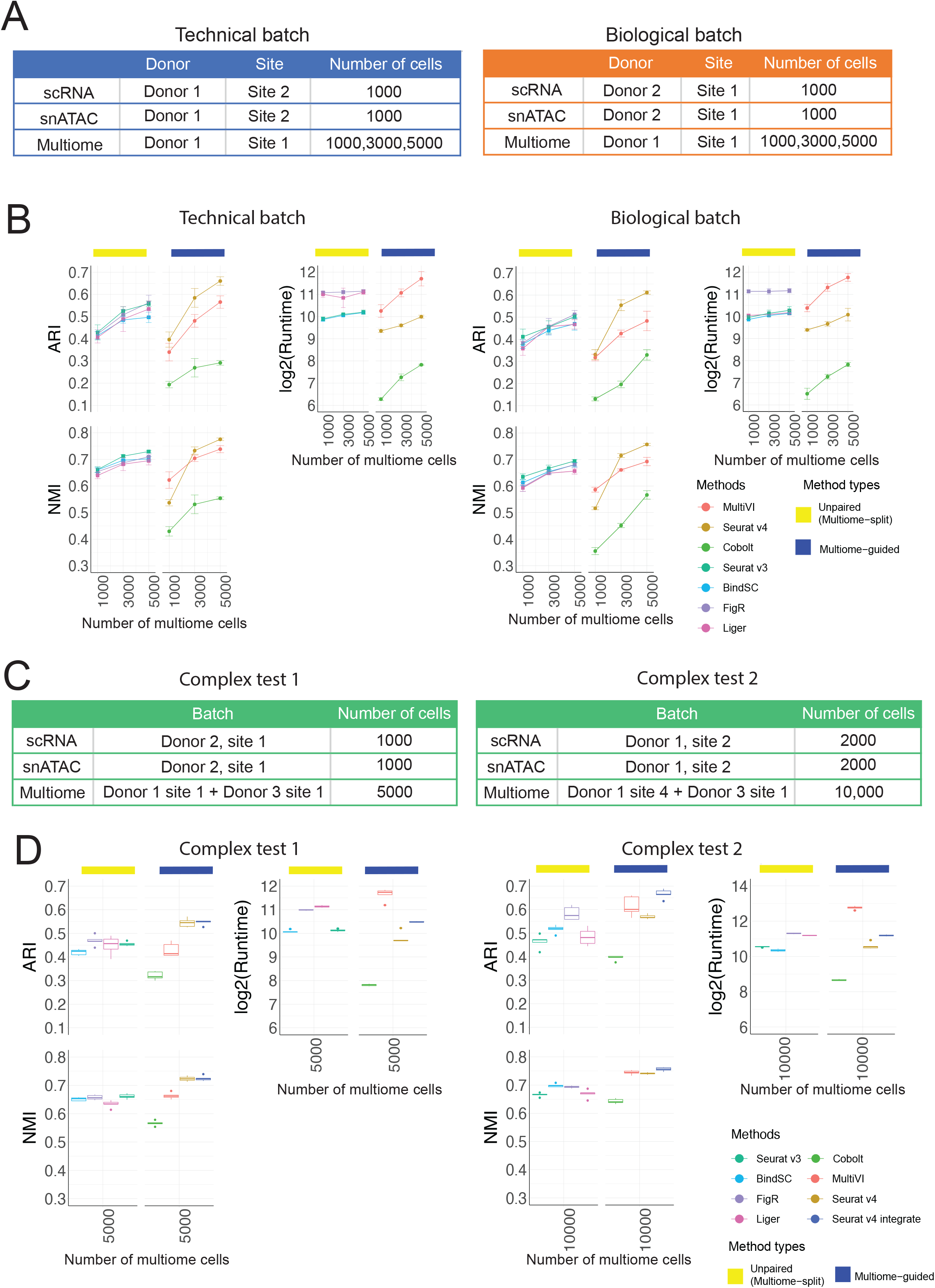
Evaluation of integration performance in the presence of batch effects. (A) Simulation details for the constructed data with technical batches and biological batches. (B) Performance of cell type annotation and runtime in the presence of technical and biological batches shown in (A). ARI and NMI measure agreement between predicted cell type and ground-truth labels. Runtime is measured in seconds, for each method, in log2 scale. Error bar is mean ± standard deviation. (C) Simulation details for two datasets with more complex batch structures. (D) Performance of cell type annotation and runtime in the presence of technical and biological batches shown in (C). ARI and NMI measure agreement between predicted cell type and ground-truth labels. Runtime is measured in seconds, for each method, in log2 scale. Whisker is 1.5 times the inter-quartile range.

To further challenge all methods in the situation of complex mixtures of samples, we considered a situation where the multiome sample includes cells from a mixture of two donors, and the scRNA-seq and snATAC-seq data come from the same or different research sites. Due to batch effects in the multiome samples, we added one more category called ‘Seurat v4 integrate’, in which the integration of samples was first done on each modality separately, then two modalities were joined using the Seurat v4 weighted nearest neighbor approach, and lastly combined with the single modality dataset (see more in Supplementary methods). Figure 4D shows that in the case of low batch effects between the two donors, Seurat v4 and ‘Seurat v4 integrate’ performed similarly well at annotating cell types. However, in the presence of stronger batch effects, ‘Seurat v4 integrate’ outperformed all other methods for cell type annotation, with much higher cell type separation as measured in cell type average silhouette width (ASW) (Supplementary Figure 13). From the UMAP projection in Supplementary Figure 14, we see that ‘Seurat v4 integrate’ mixes cells from the two multiome samples much better than Seurat v4.

Therefore, when the multiome data include two donors with strong batch effects, integration across the batches is required before mapping the single-modality datasets.

## Discussion

In summary, we evaluated seven multi-omic integration methods under three realistic scenarios. Firstly, we showed that the incorporation of multiome data improves the cell type annotation accuracy of scRNA-seq and snATAC-seq data when there are sufficient number of cells in the multiome data to reveal cell type identities. Secondly, we showed that the number of cells in the multiome data plays a more important role than sequencing depth per cell for cell type annotation accuracy. Thus, when generating a multiome dataset with a fixed budget, a better strategy is to profile more cells so that rare cell types can be captured. Lastly, when the three datasets to be integrated are confounded by batch effects, Seurat v4 resulted in the best cell type annotation accuracy.

In all evaluations, Seurat v4 demonstrated superior performance at resolving cell type heterogeneity when data from many multiome-profiled cells are available. This makes sense as Seurat v4 is a supervised approach in which single-modality cells are merely projected to the integrated space learned from the multiome dataset. Therefore, when the multiome data have an insufficient number of cells to reveal accurate cell types, the integration will lead to poor annotation accuracy. The other two multiome-guided methods, e.g., Cobolt and MultiVI, claim to be able to make use of all three data types. The hope is that the single-modality cells can help the clustering when multiome cells are small. However, as shown in Figures 2 and 3, both Cobolt and MultiVI performed worse than the unpaired integration methods that do not leverage the paired relationship of the multiome data. Therefore, when the multiome dataset has a small number of cells, it is better to treat the multiome cells as unpaired and append them to the single-modality datasets for the integration of three datasets.

There are several limitations of this study. Firstly, our simulations represent the most ideal situation, where the single-modality cells are generated from the exact same dataset as the multiome cells. In reality, the single-modality and the multiome data are generated from different experimental kits that could have slight differences since the multiome workflow is optimized to capture both gene expression and chromatin accessibility. Moreover, the gene expression captured through the multiome workflow is in fact measuring mRNA in individual nuclei, while scRNA-seq captures mRNA in whole cells. Slight differences between snRNA-seq and scRNA-seq datasets have been reported [19]. Lastly, the PBMC dataset did not have expert-annotated cell type labels. We followed a tutorial by Seurat v4 to obtain annotations [20], thus the evaluation of PBMC-simulated scenarios might favor Seurat v4. However, the BMMC data were manually annotated by experts and Seurat v4 still showed outstanding performance in evaluations based on this dataset.

Secondly, although Seurat v4 was the best at annotating cell types, it performed worse than unpaired integration methods at recovering peak-gene associations. Furthermore, even the best method only revealed 45% of peak-gene pairs detected in the paired multiome dataset, and many of the detected pairs are false positives. Moreover, we did not explore the possibility of imputing chromatin accessibility from scRNA-seq or appending imputed profile with observed multiome sample. To truly integrate the three data types and understand the underlying *cis*-regulatory logic, one would hope to impute the missing modality for both the scRNA-seq and snATAC-seq data, and then append the imputed profiles with the multiome dataset to identify peak-gene pairs with the largest number of cells. Therefore, additional work needs to be done to evaluate the performance of different methods in jointly integrating the imputed single-modality datasets with the multiome samples for downstream analyses.

### Conclusions

Our benchmarking evaluations showed that multiome data are helpful for annotating single-modality data. The number of cells in the multiome data is critical to ensure a good cell type annotation after integration and the exact number of cells depends on the complexity of the biological system. When generating a multiome dataset, the number of cells is more important than sequencing depth for cell type annotation. Lastly, Seurat v4 is the best at integrating scRNA-seq, snATAC-seq, and multiome data even in the presence of complex batch effects.

## Methods

### Datasets

#### Peripheral blood mononuclear cell (PBMC) dataset

This dataset was generated using the 10x Genomics Single Cell Multiome ATAC + Gene Expression kit [10]. The PBMC dataset with granulocytes removed was downloaded from the 10x Genomics website, which included 11,909 cells. The dataset was processed and annotated into 30 cell types following the Seurat tutorial [7, 20]. We grouped similar cell types and refined the annotations into 9 broad cell types (similar to the level 1 categories from the Azimuth database [3]): B-cells (‘B’), CD4 T cells (‘CD4 T’), CD8 Naïve T cells (‘CD8 Naïve’), CD8 Effector T cells (‘CD8 TEM’), Dendritic cells (‘DC’), Monocytes (‘Mono’), Nature killer cell (‘NK’), other T cell (‘other_T’), and other cell categories (‘other’). The ATAC-seq profile released on 10x Genomics website was counting the Tn5 insertion events in each genomic region. Here, we retabulated the cell-peak matrix by the number of reads overlapping each genomic region, using the Signac’s FeatureMatrix function [21]. We used the peak-based counting result as input for the peak-gene pair identification (described below) and subsequent simulations. The list of peak-gene pairs identified using all cells in the multiome dataset (10,412 cells) is treated as the ground truth when calculating percentage of peak-gene pair recovery or F1 score. ‘Other_T’ and ‘other’ cells were excluded from the data simulation due to their extensive separation in the UMAP embedding. After removal of cells, there are 10,085 cells used for simulation.

#### Bone marrow mononuclear cells (BMMC) dataset

This dataset was generated as part of the “Open Problems in Single-cell Analysis” competition [12]. BMMC cells from nine healthy donors were profiled at four different research sites using the 10x Multiome ATAC + Gene Expression kit. The dataset was analyzed by Lance and colleagues [12], who annotated the cells into 22 cell types. The values in the cell-peak matrix of the ATAC-seq data was also the insertion-based counting, so we again converted it into peak-based counting as mentioned above. Data simulations related to Figures 2 and 3 were performed using cells from the site 1 donor 2 (S1D2) BMMC sample. This sample contains 6,740 cells, annotated into 21 cell types. The peak-gene pair prediction accuracies shown in Figures 2 and 3 were calculated by comparing the result to a ground-truth list generated with the S1D2 sample. To simulate technical batch and biological batch effects (Figure 4), we used cells generated at research site 1 or from donor 1, which includes a total of 29,486 cells, composed of 21 cell types (Supplementary Figure 1B).

#### Evaluation metrics

##### Annotation accuracy

Each integration method returns an integrated latent embedding matrix for cells. Louvain clustering was performed to identify k clusters, in which k is the number of cell types in the ground-truth annotation. To evaluate annotation accuracy, Adjusted Rand Index (ARI) [14] and Normalized Mutual Information (NMI) [15] from the Scib package (v1.0.2) [18] were calculated to compare the predicted cluster labels with the ground truth. Specifically, ARI compares every pair of cells in the dataset and calculates a similarity measurement by considering the number of cell pairs that are in the same cluster in both annotation results, versus the number of cell pairs showing discordant annotations. This metric is then adjusted by chance, as there will be a non-zero similarity between the two clustering results just due to random permutation of labels. The resulting metric ranges from 0 to 1 in which 1 means perfect matching between the two results while 0 means random labeling of cells. NMI is another measurement commonly used for comparison of two clustering results. NMI measures if knowing one label provides information about the other label. If the two lists are highly correlated, then it has high mutual information. NMI is then normalized by a factor to control for differences due to the number of clusters in each set of labels.

##### Cell type separation

We evaluated the separation of clusters and the tightness of cells in the integrated latent space derived from each method. We calculated cell type-specific average silhouette width (ASW) [18], using the ground-truth annotation and the joint embeddings. The resulting score is between 0 and 1 in which 1 means small intra-cluster distance and high inter-cluster distance. We also calculated a cell type Local Inverse Simpson’s Index (cLISI) [18], which is an adaptation of LISI previously used to quantify the degree of batch effects [17]. Here, cLISI was calculated using the ground-truth labels again in which it evaluates how many cells need to be drawn from a cell’s neighborhood to draw a second cell of the same type. The score is normalized again so that 1 means good local neighborhood preservation of the same cell type while 0 is otherwise.

##### Batch mixing

To evaluate batch mixing, two metrics were employed. A batch ASW score was used to evaluate the within-batch distance and the across-batch distance [18]. The score was rescaled so that 0 is the worst and 1 is the best separation. To evaluate the local neighborhood accuracy, k-nearest neighbor batch effect test (kBET) was also performed [16]. Specifically, kBET measures the difference between observed batch frequency in the k-nearest neighbors compared to an expected frequency based on the number of cells in each batch. The value is rescaled to 0 and 1 in which 1 represents the optimal mixing of cells from different batches in which cells in the neighborhood are highly similar to the expected frequency.

##### Peak-gene pair recovery

To identify correlated peak-gene pairs, we used the methodology introduced in the SHARE-seq paper [1]. Specifically, a Pearson correlation is calculated between the raw accessibility count of every peak and the normalized UMI count of every gene if the peak is within 50,000 base pairs from the transcription start site (TSS) of the gene. The null distribution of correlation coefficients was then generated through selecting 100 peaks that have similar GC content, length, and accessibility as the target peak, and calculating correlation of the background peaks and the target gene. A one-sided t-test was used to calculate a p-value for every peak-gene pair by comparing to the background peaks and the peak-gene pairs with p-value less than 0.05 and z-score greater than 0.05 identified as significant peak-gene pairs. Associated peak-gene pairs were identified using all cells from each dataset. To evaluate the performance of each method at imputing gene expression from snATAC-seq data, a peak-gene association was calculated in the same manner using the raw cell-peak count of the unpaired ATAC data and the predicted gene expression generated by the evaluated methods. To evaluate the *in silico* imputed gene expression results, we calculated the percentage of peak-gene pairs recovered using the imputed gene expression and the observed snATAC-seq peak counts. To account for false negative results, we calculated an F1 score. Thus, the peak-gene pair percent recovery and the F1 score were used to evaluate each method that can impute missing gene expression.

#### Evaluation scenarios

We simulated three scenarios to evaluate the performance of each method. For each scenario, we simulated five independent replicates. Details regarding how each method was implemented are described in the Supplementary Methods.

#### Scenario 1: evaluating the effect of multiome cells on single-modality integration

##### Data simulation

In this task, we first defined the number of cells to be drawn for each data type with an example shown in Figure 2A. Then, we randomly selected cells from the ground-truth multiome dataset according to the desired number of cells for each data type. For scRNA-seq, we kept the gene expression matrix; for snATAC-seq, we kept the cell-by-peak matrix and the fragment file; lastly, for the multiome sample, we kept all three data files. The cells were sampled without replacement.

##### Evaluated methods

We first ran the four unpaired integration methods (Seurat v3, LIGER, FigR, and bindSC) to integrate the simulated scRNA-seq and snATAC-seq datasets and the results were summarized under the ‘Unpaired’ categories. To make use of the multiome data, we ran the four methods again, with the multiome cells treated as unpaired. Specifically, the RNA profile from the multiome cells was appended to the scRNA-seq dataset, and the ATAC-seq profile was appended to the snATAC-seq dataset. The results from this category were summarized under ‘Unpaired (multiome-split)’. Lastly, we ran the multiome-guided methods with the scRNA-seq, snATAC-seq, and multiome datasets as input.

##### Evaluations

To evaluate if the presence of multiome cells improves the integration of single-modality datasets, we evaluated the annotation accuracy, peak-gene pair recovery, cell type separation, and batch mixing of the scRNA-seq and snATAC-seq cells.

#### Scenario 2: evaluating the impact of sequencing depth in multiome cells on multi-omic data integration

##### Data simulation

For this task, we first defined the number of cells in each data type as well as the percentage of original depth the multiome cells will be down-sampled to; an example is shown in Figure 3A. We first generated the three data types according to the number of cells defined. Then, we performed depth-down-sampling for both the gene expression and chromatin accessibility profiles of the multiome dataset. To down-sample the cell-by-gene count matrix for gene expression, we used Scuttle::downsample [22] to reduce the sample depth to a percentage of the original dataset. To down-sample the ATAC-seq depth, we performed down-sampling on the fragment file and then regenerated the cell-by-peak count matrix. Specifically, we first counted the number of fragments corresponding to the selected cells, then we calculated the target depth by multiplying the original depth to the percentage factor. We randomly selected the number of reads as calculated, without replacement, and saved this file as the new fragment file. Then the down-sampled fragment file was sorted, recompressed, indexed with tabix and, tabulated into peak counts with the original feature set with Signac:: FeatureMatrix [21] function. This often resulted in less reduction in peak counts, as some of the fragments removed were not previously assigned to the peaks.

##### Evaluated methods

We ran the unpaired integration methods with the multiome data appended to the single-modality datasets as described above, the results were summarized under ‘Unpaired (multiome-split)’. We also ran the three multiome-guided methods.

##### Evaluations

The evaluation of annotation accuracy, cell type separation and batch mixing were calculated using all cells present in simulated scRNA-seq, snATAC-seq, and the multiome datasets. Given how the multiome data were split and appended to the single-modality datasets for the ‘unpaired (multiome-split)’ category, it resulted in doubling the number of multiome cells. Thus, to ensure a fair comparison between the two categories of methods, half of the multiome cells appended to the RNA-seq were dropped while the other half of the multiome cells appended to the ATAC-seq were dropped. As a result, the same number of cells was evaluated for the ‘unpaired (multiome-split)’ and ‘multiome-guided’ methods.

#### Scenario 3: evaluating the impact of batch effects on multi-omic data integration

##### Data simulation

The analysis of batch effects was only possible for the BMMC dataset. As mentioned before, the BMMC dataset contains multiome cells generated at four different research sites and nine donors. To create different types of batches, we used the multiome cells from donor 1 but processed at three different sites (S1D1, S2D1, S4D1) as the data source to generate technical batches. We used the multiome cells generated at research site 1 but from different donors (S1D1, S1D2, S1D3) as the source of biological batches. To generate scenarios with mixed technical and biological batch effects, we created more complex batch structures described as ‘complex test’ in Figure 4D using all samples that were either generated at research site 1 or donor 1. After defining which sample each data type comes from and the number of cells, the simulation is the same as described in ‘Sceanrio 1’, in which cells were randomly drawn from the ground-truth multiome dataset to simulate scRNA-seq, snATAC-seq, and multiome samples.

##### Evaluated methods

The same seven methods, four from the ‘unpaired (multiome-split)’ and three from ‘multiome-guided’ were ran. For situations were multiome were composed of two donors, an additional variation of Seurat v4 was added, termed ‘Seurat v4 integrate’. Specifically, the two multiome datasets were first integrated across donors to generate one integrated reference before it was used to integrate scRNA-seq and snATAC-seq datasets.

##### Evaluations

We calculated metrics measuring annotation accuracy, cell type separation, and batch mixing. For batch mixing, we calculated both the mixing of data types, as well as the mixing of samples. Similar to what was described in ‘Scenario 2’, to ensure that the same number of cells were evaluated for the unpaired (multiome-split) methods and the multiom-guided methods, half of multiome cells appended to the RNA-seq and the other half of the ATAC-seq dataset were dropped.

## Declarations

### Availability of data and materials

The source codes for simulation and evaluations are available online on GitHub at https://github.com/myylee/benchmark_sc_multiomic_integration. For the multiome datasets used to generate simulated data, the 10x PBMC dataset was downloaded from https://www.genomics.com/resources/datasets/pbmc-from-a-healthy-donor-granulocytes-removed-through-cell-sorting-10-k-1-standard-2-0-0 and the BMMC dataset was downloaded from GEO accession GSE194122, and the fragment files were obtained from the authors [23].

## Authors’ contributions

M.Y.Y.L., M.L., K.H.K. conceived this project and designed the framework together. M.Y.Y.L. performed the simulations and evaluations with guidance from M.L. All authors wrote and edited the final manuscript. M.L. and K.H.K. supervised the study.

## Acknowledgements

We thank everyone in the Li Lab and Kaestner lab for helpful discussions, especially Dr. Elisabetta Manduchi and Dr. Jonathan Schug.

