## Supplementary materials for "Benchmarking algorithms for joint integration of unpaired and paired single-cell RNA-seq and ATAC-seq data"

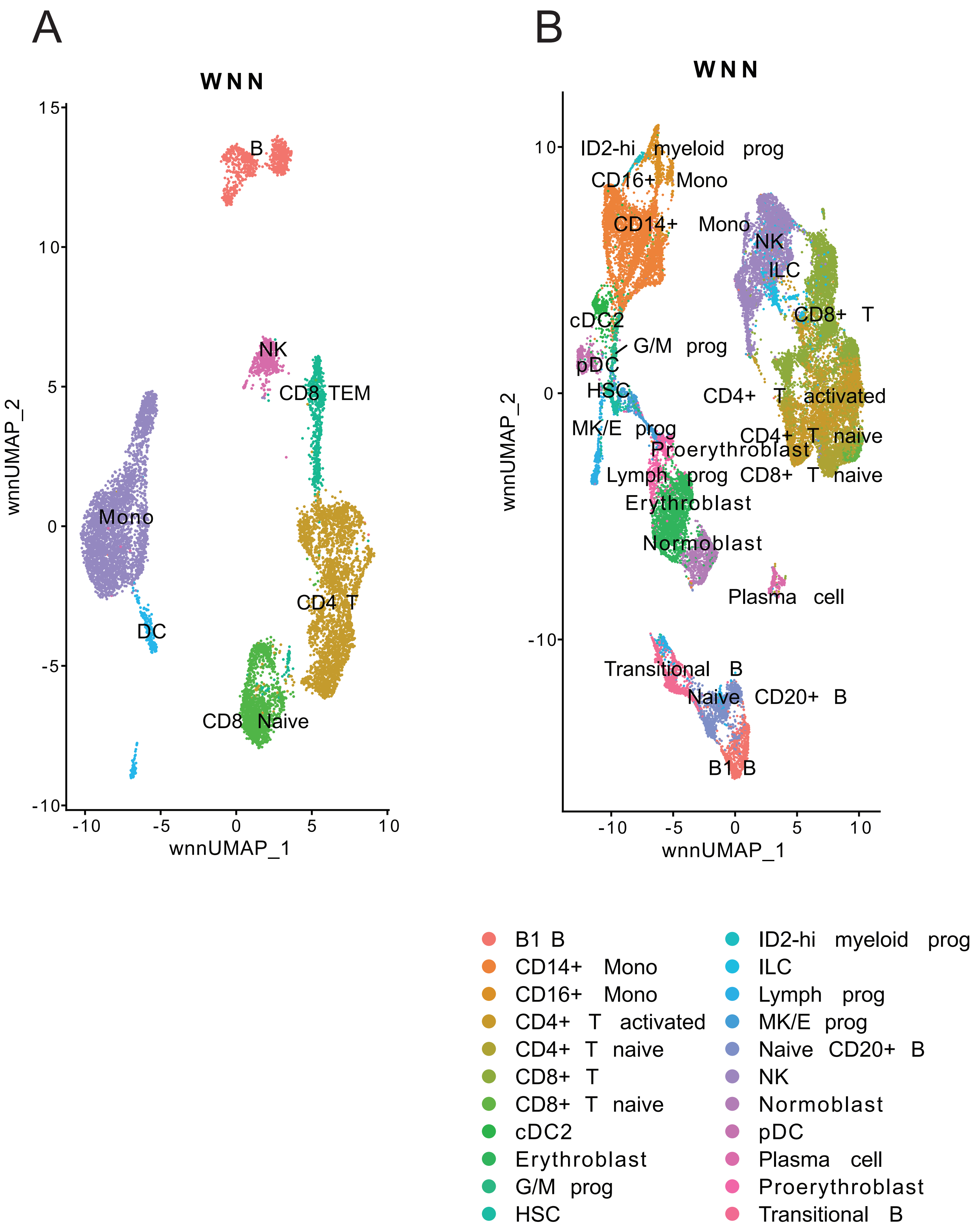


Supplementary Figure 1: UMAP plots colored by cell types. (A) PBMC-based simulations. (B) BMMC-based simulations.

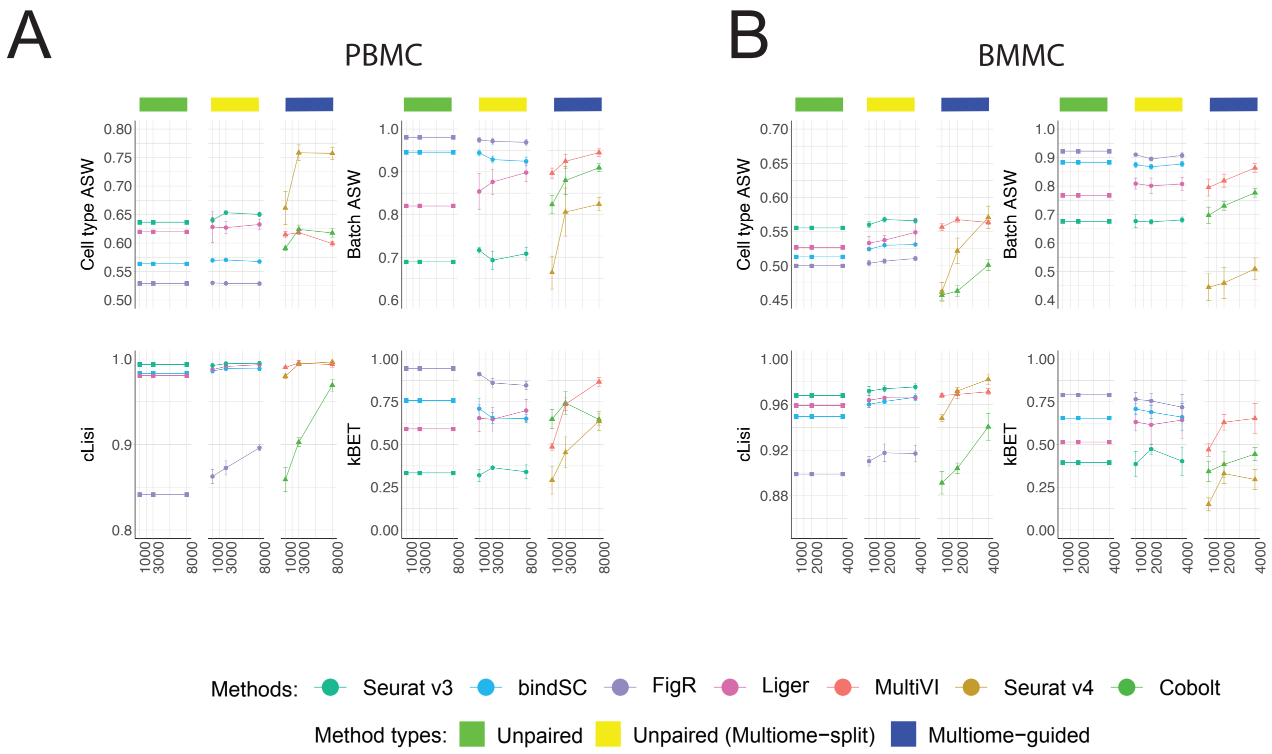


Supplementary Figure 2: Additional evaluation metrics for each method at integrating single-modality cells in the presence of multiome data, as described in Figure 2. (A) PBMC-based simulations. (B) BMMC-based simulations. Cell type average silhouette width (ASW) and cell type Local Inverse Simpson's Index (cLISI) measure separation of cell types. Batch ASW and k-nearest neighbor batch effect test (kBET) measure the mixing of scRNA-seq and snATAC-seq cells. Error bar is mean ± standard deviation.

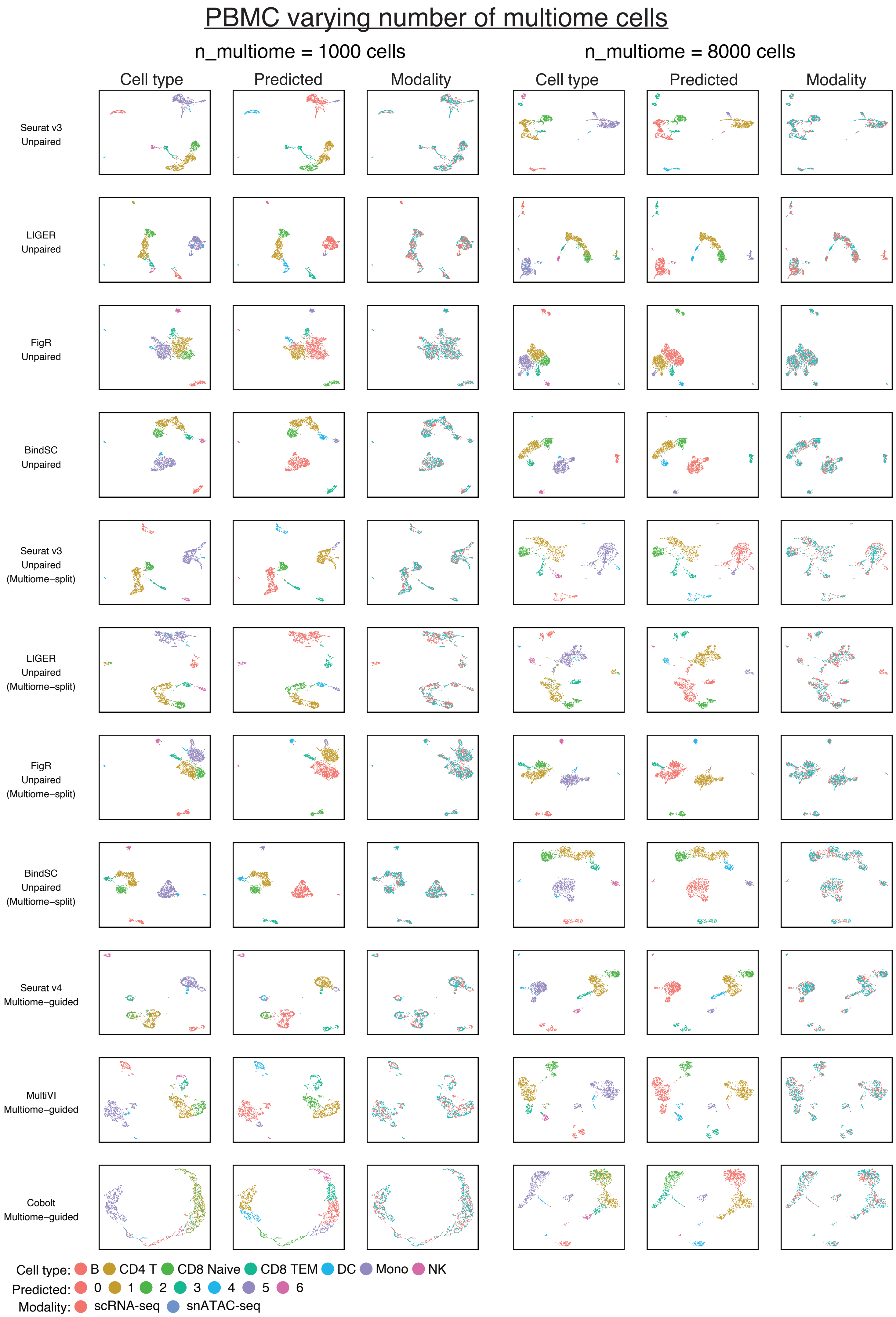


Supplementary Figure 3: UMAP plots for the PBMC-based simulations shown in Figure 2B.

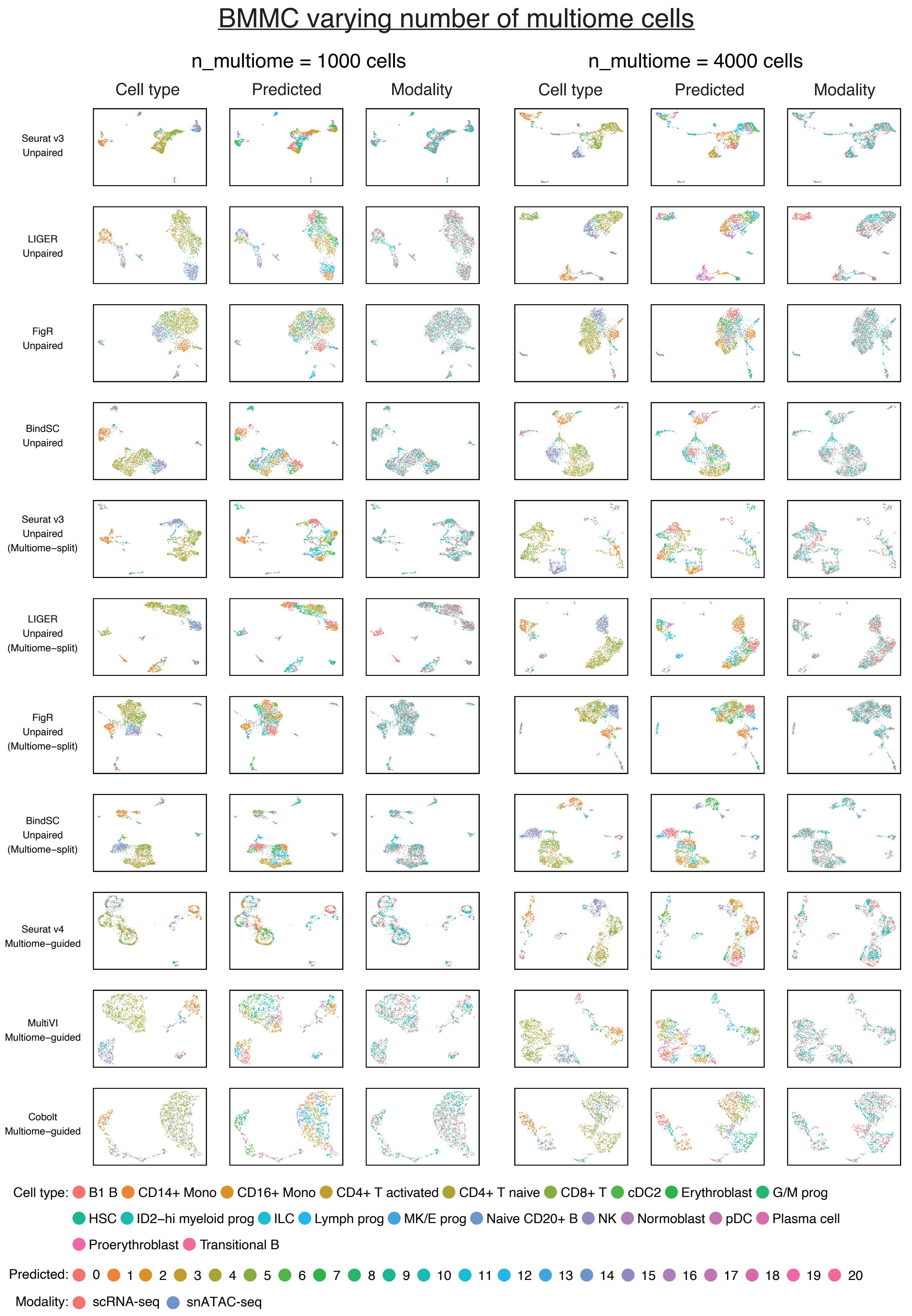


Supplementary Figure 4: UMAP plots for the BMMC-based simulations shown in Figure 2C.

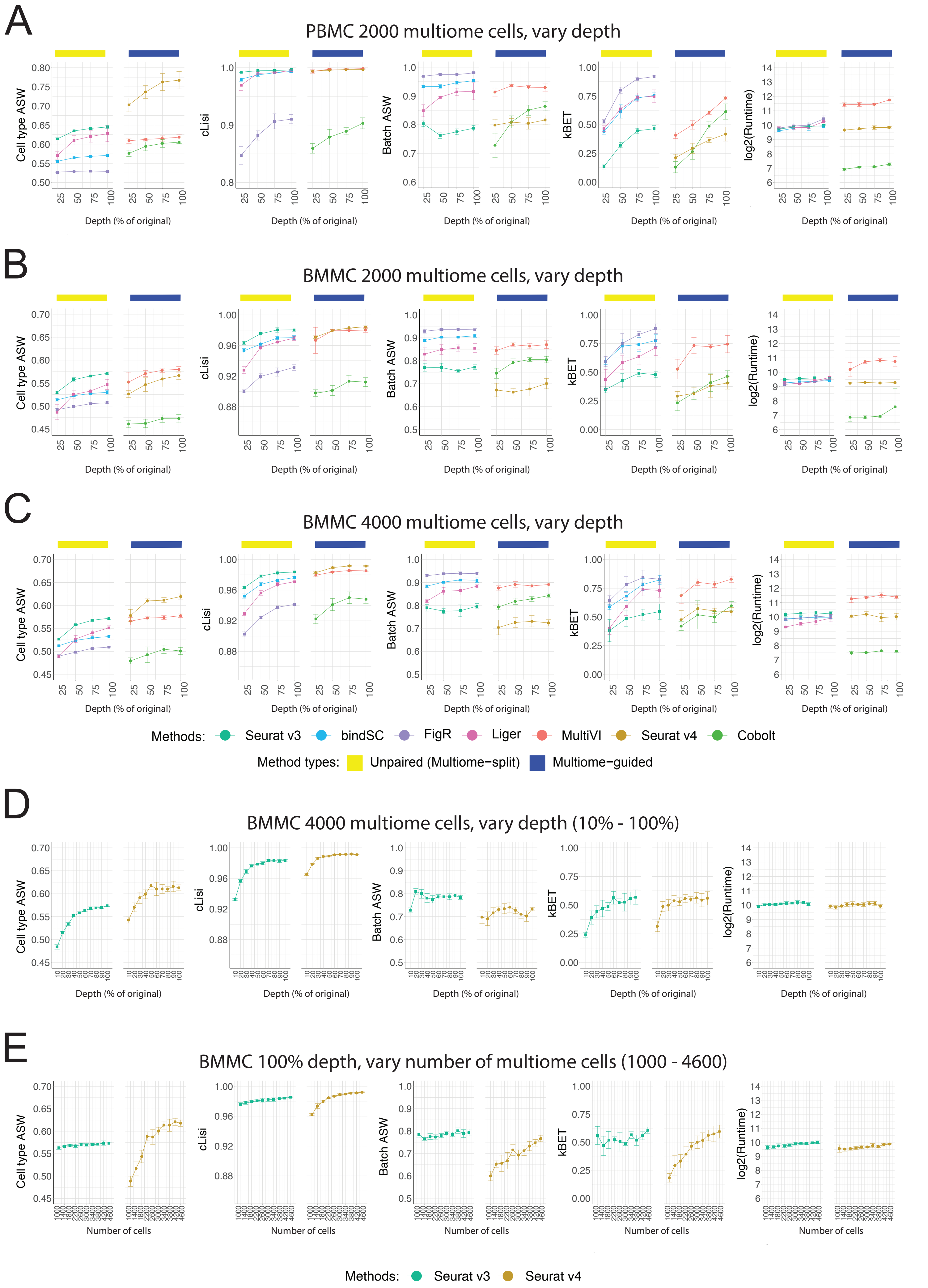

A
PBMC 2000 multiome cells, vary depth
B
BMMC 2000 multiome cells, vary depth
C
BMMC 4000 multiome cells, vary depth
Cell type ASW
cLisi
Batch ASW
kBET
log2(Runtime)
Depth (% of original)
Methods: Seurat v3 bindSC FigR Liger MultiVI Seurat v4 Cobolt
Method types: Unpaired (Multiome−split) Multiome−guided
D
BMMC 4000 multiome cells, vary depth (10% - 100%)
E
BMMC 100% depth, vary number of multiome cells (1000 - 4600)
Number of cells
Methods: Seurat v3 Seurat v4

**Supplementary Figure 5:** Additional evaluation metrics for each method at integrating scRNA-seq, snATAC-seq and multiome data, at different depths of the multiome data, as described in Figure 3. (A) PBMC-based simulations. (B) BMMC-based simulations with 2000 multiome cells. (C) BMMC-based simulations with 4000 multiome cells. (D) BMMC-based simulations with 4000 cells at 10 different depths. (E) BMMC-based simulations at 100% of depth but with 10 different numbers of multiome cells. Cell type ASW and cLISI measure separation of cell types. Batch ASW and kBET measure the mixing of scRNA-seq, snATAC-seq, and multiome cells. Runtime is measured in seconds, for each method, in log2 scale. Error bar is mean ± standard deviation.

### PBMC varying depth of multiome cells

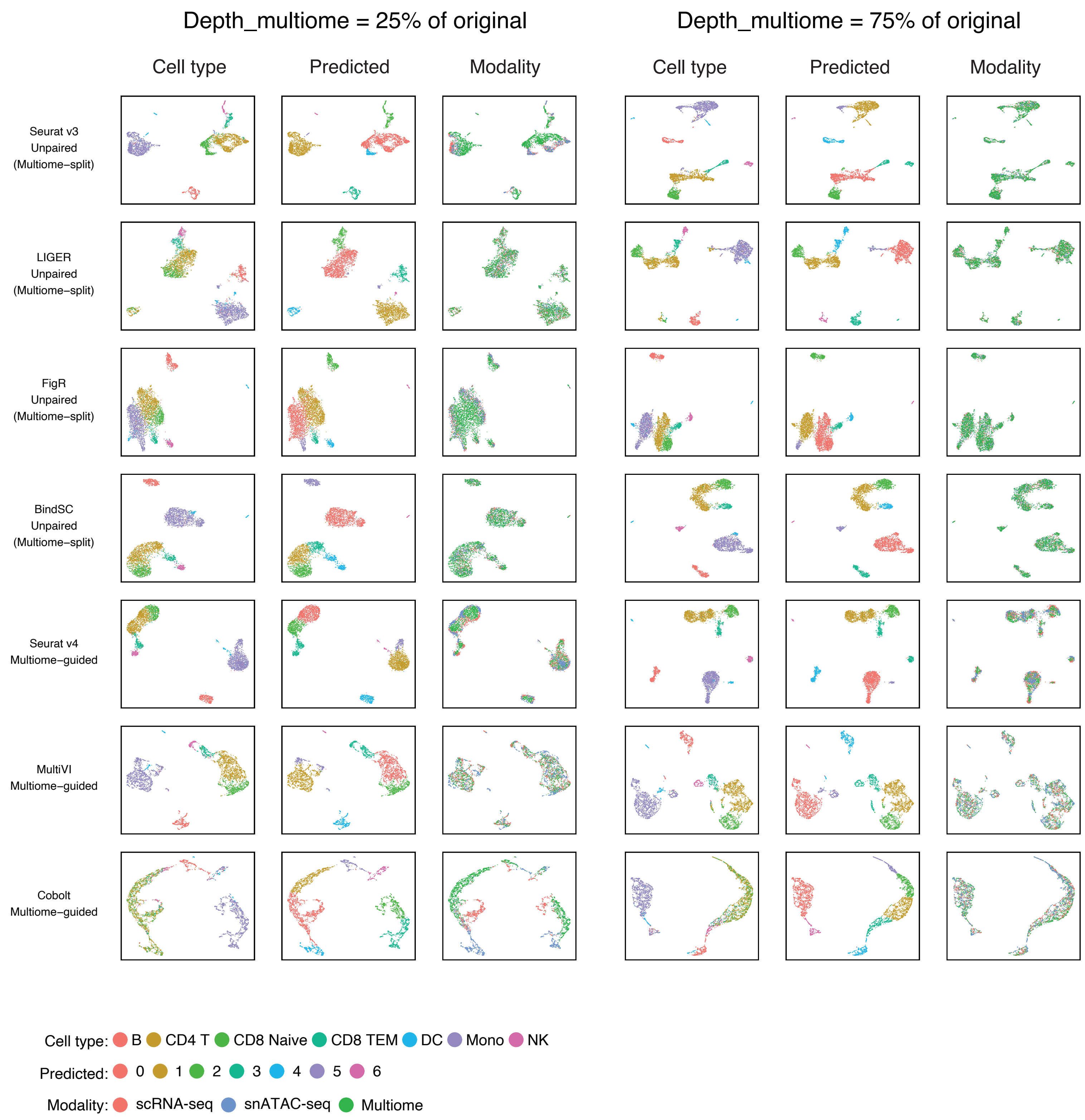


Supplementary Figure 6: UMAP plots for the PBMC-based simulations shown in Figure 3B.

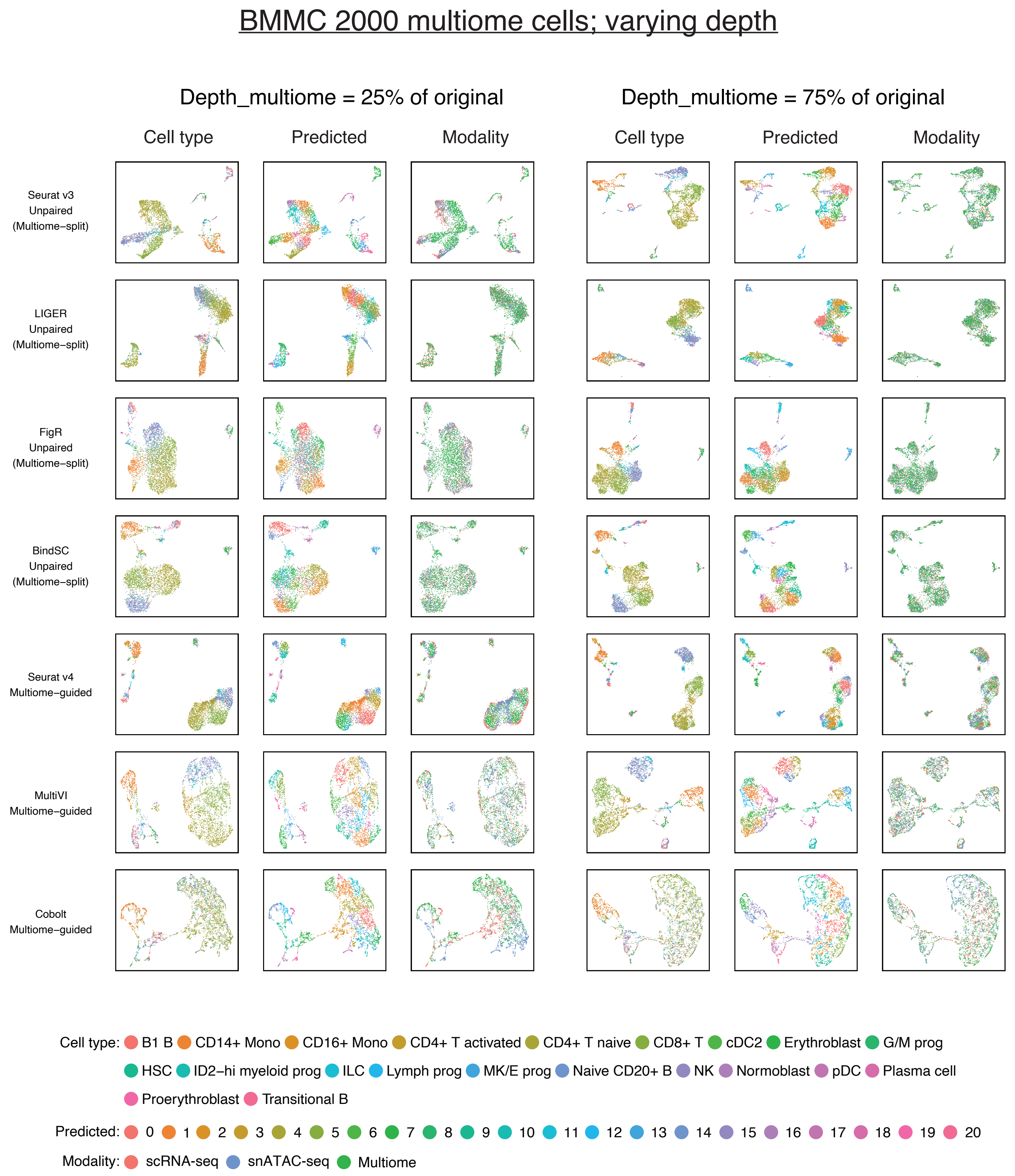


Supplementary Figure 7: UMAP plots for the BMMC-based simulations with 2000 multiome cells shown in Figure 3C (left).

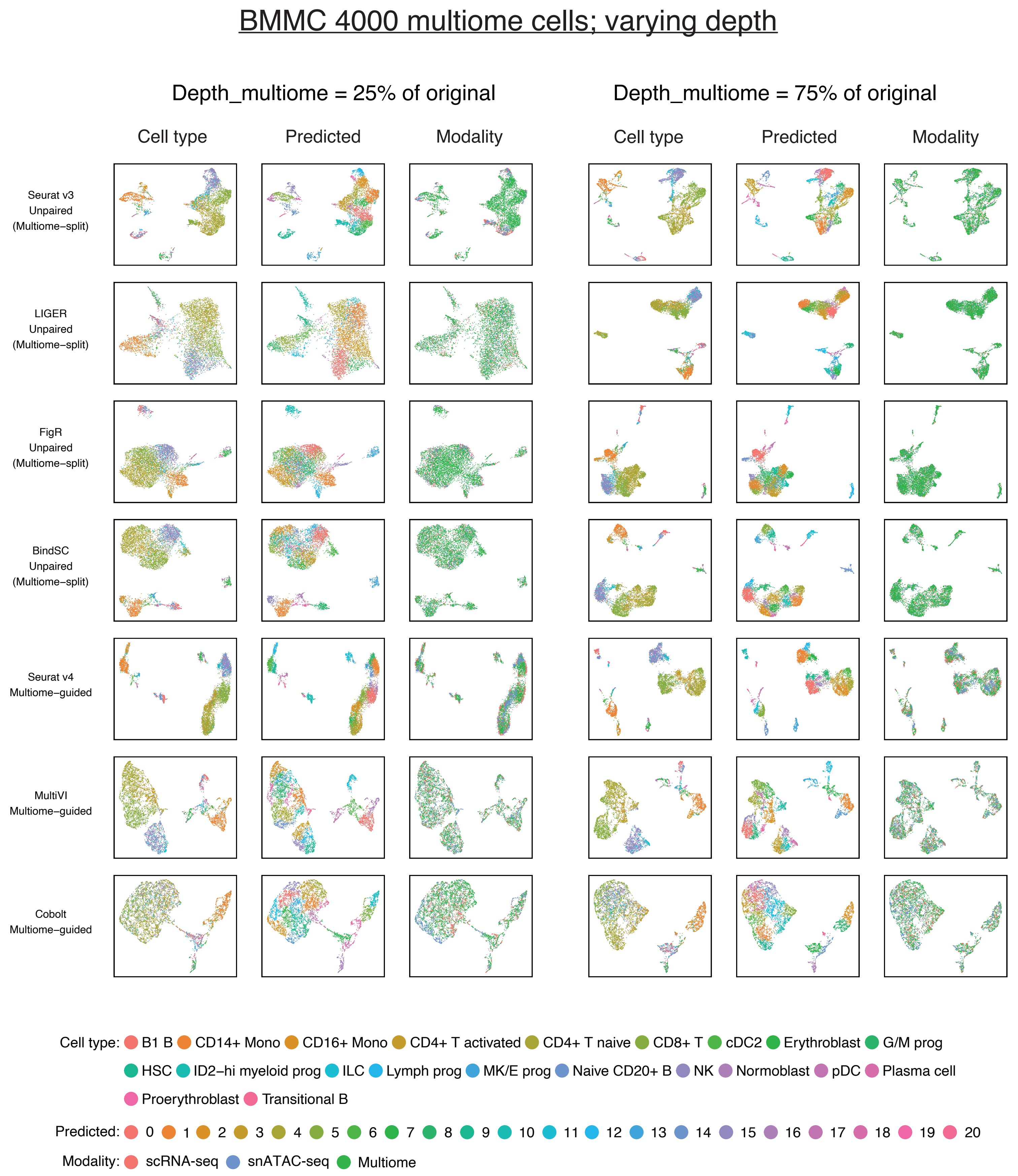


Supplementary Figure 8: UMAP plots for the BMMC-based simulations with 4000 multiome cells shown in Figure 3C (right).

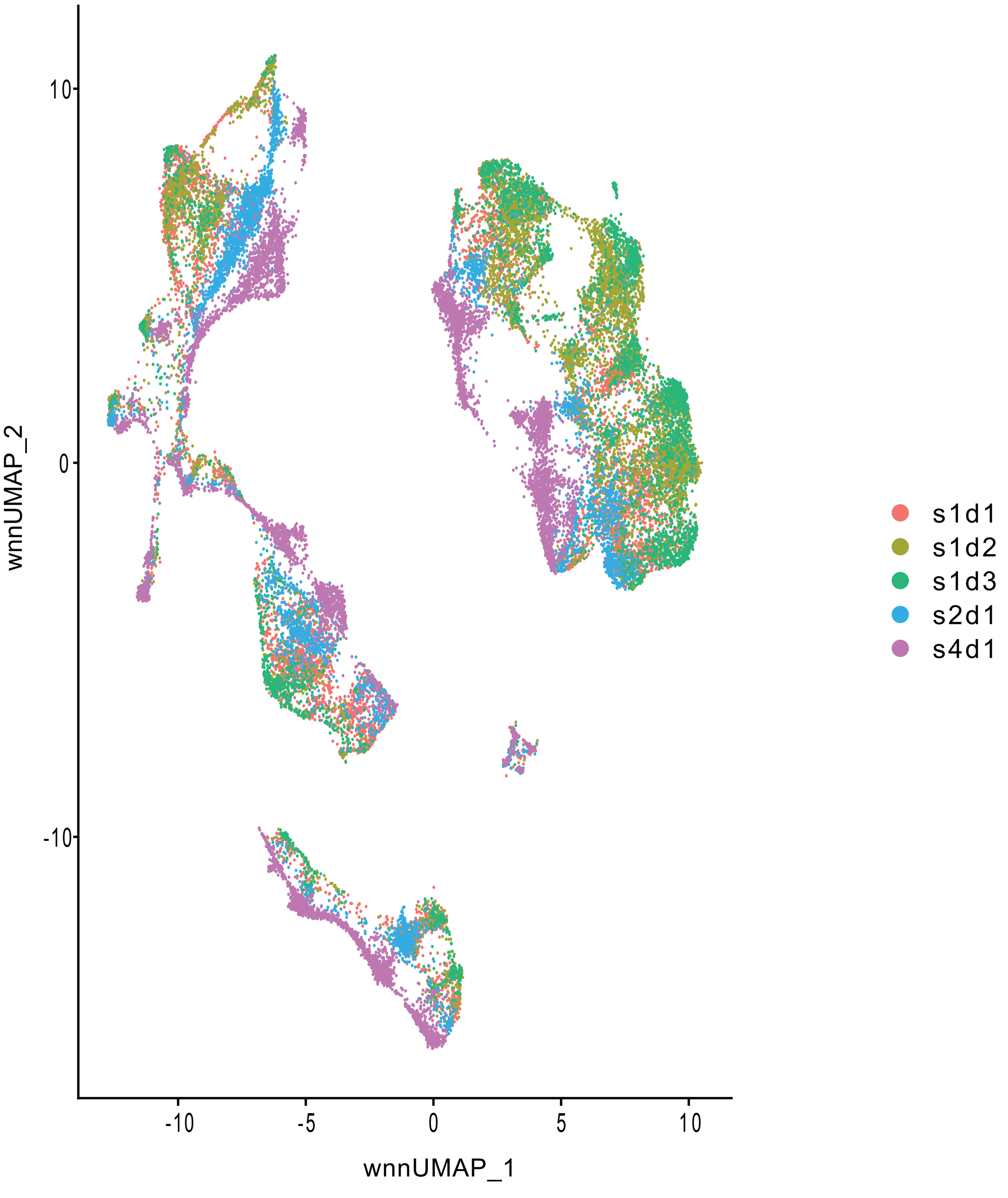


Supplementary Figure 9: UMAP plots colored by sample origins, containing 5 BMMC samples generated from research site 1 or donor 1 samples.

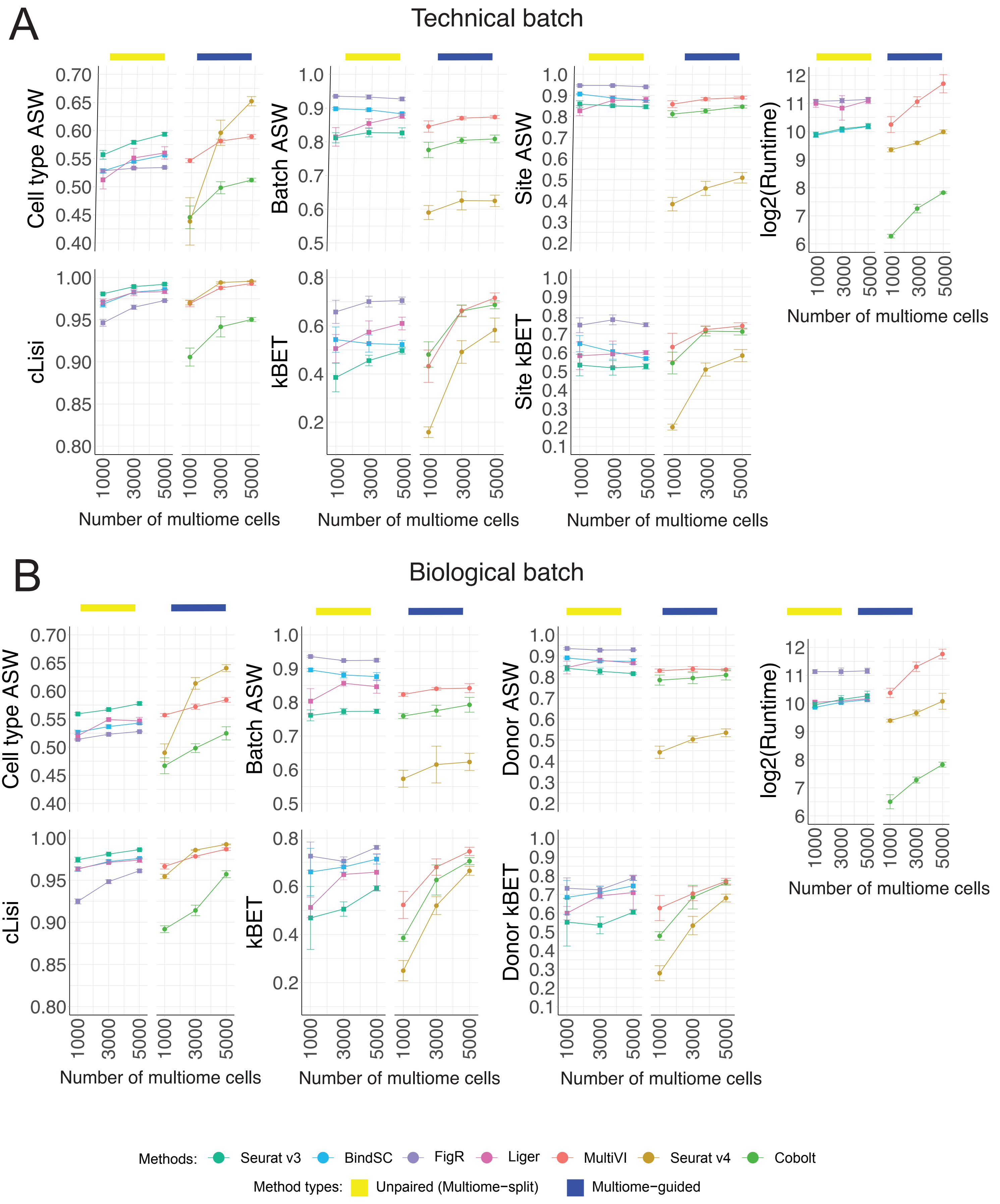


Supplementary Figure 10: Additional evaluation metrics for each method at integrating scRNA-seq, snATAC-seq and multiome data, in the presence of batch effects as described in Figure 4B. (A) Technical batch effect (Figure 4B left). (B) Biological batch effect (Figure 4B right). Cell type ASW and cLISI measure separation of cell types. Batch ASW and batch kBET measure the mixing of scRNA-seq, snATAC-seq, and multiome cells. Site ASW and site kBET measure the mixing of cells by site ID. Donor ASW and donor kBET measure the mixing of cells by donor ID. Runtime is measured in seconds, for each method, in log2 scale. Error bar is mean ± standard deviation.

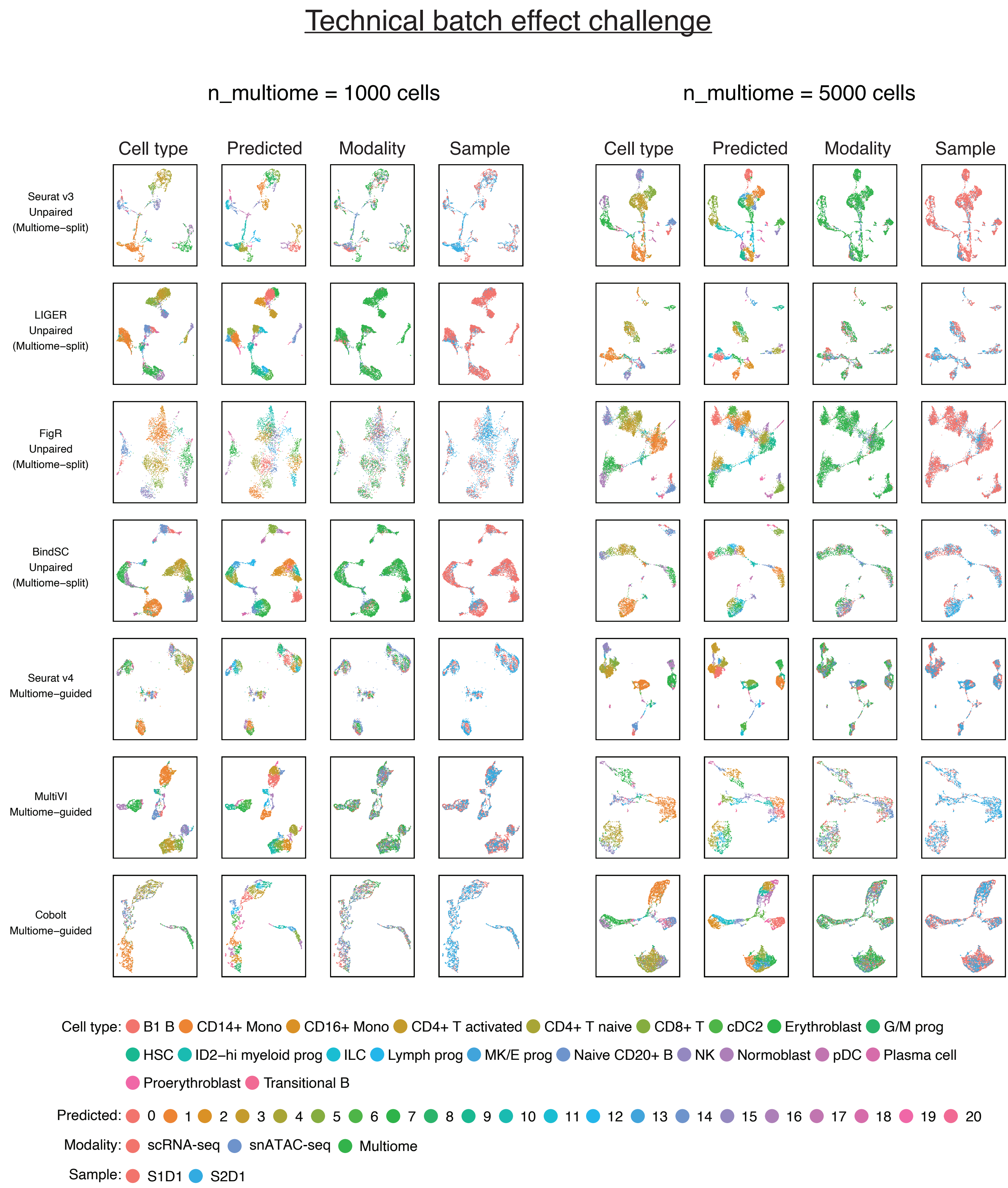


Supplementary Figure 11: UMAP plots for the BMMC-based simulations with technical batch effect challenge shown in Figure 4B (left).

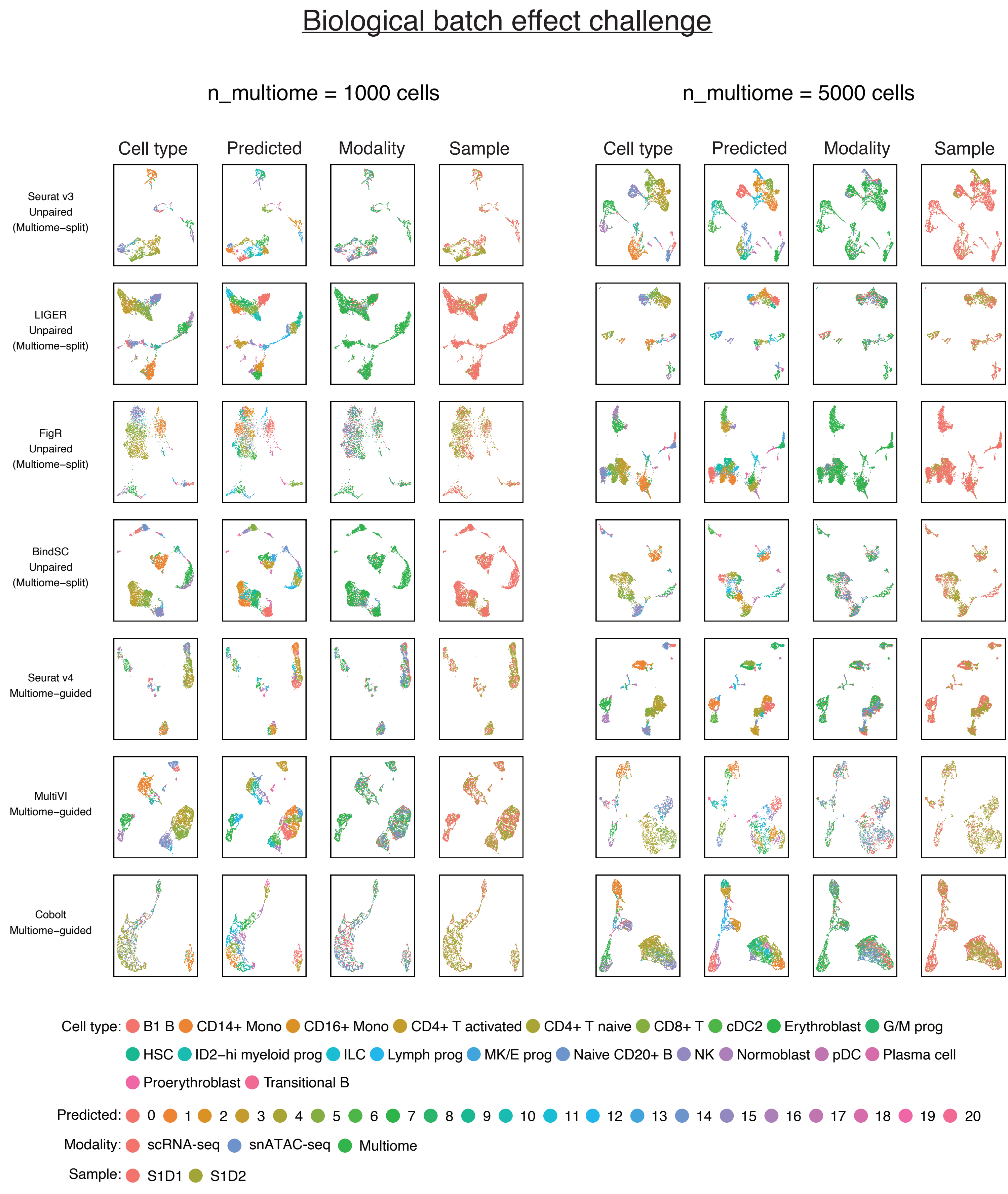


Supplementary Figure 12: UMAP plots for the BMMC-based simulations with biological batch effect challenge shown in Figure 4B (right).

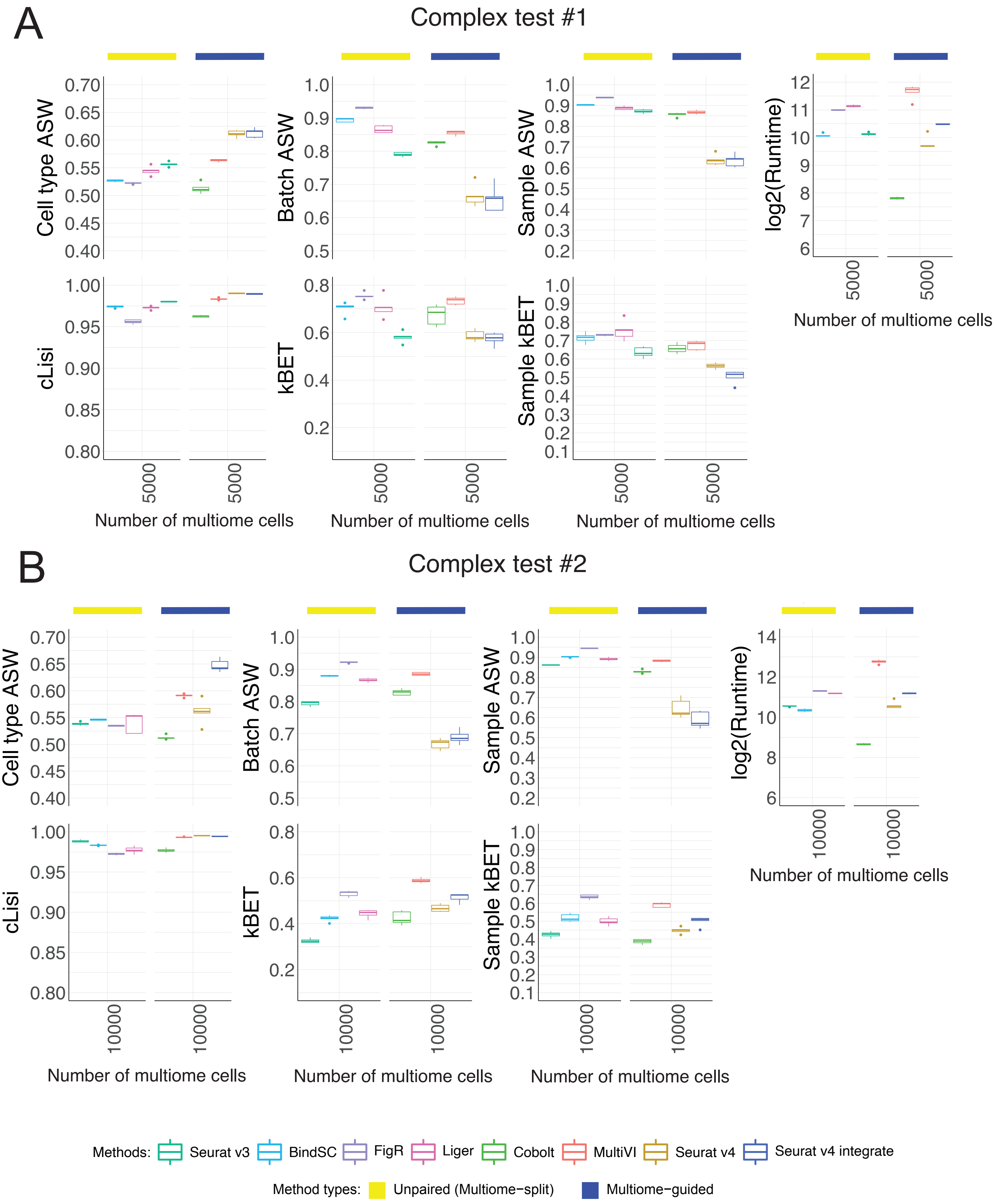


Supplementary Figure 13: Additional evaluation metrics for each method at integrating scRNA-seq, snATAC-seq and multiome data in the presence of more complex batch effects as described in Figure 4D. (A) Complex test #1 (Figure 4D left). (B) Complex test #2 (Figure 4D right). Cell type ASW and cLISI measure separation of cell types. Batch ASW and batch kBET measure the mixing of scRNA-seq, snATAC-seq, and multiome cells. Sample ASW and sample kBET measure the mixing of cells by sample ID (defined by site ID and donor ID). Runtime is measured in seconds, for each method, in log2 scale. Whisker is 1.5 times the inter-quartile range.

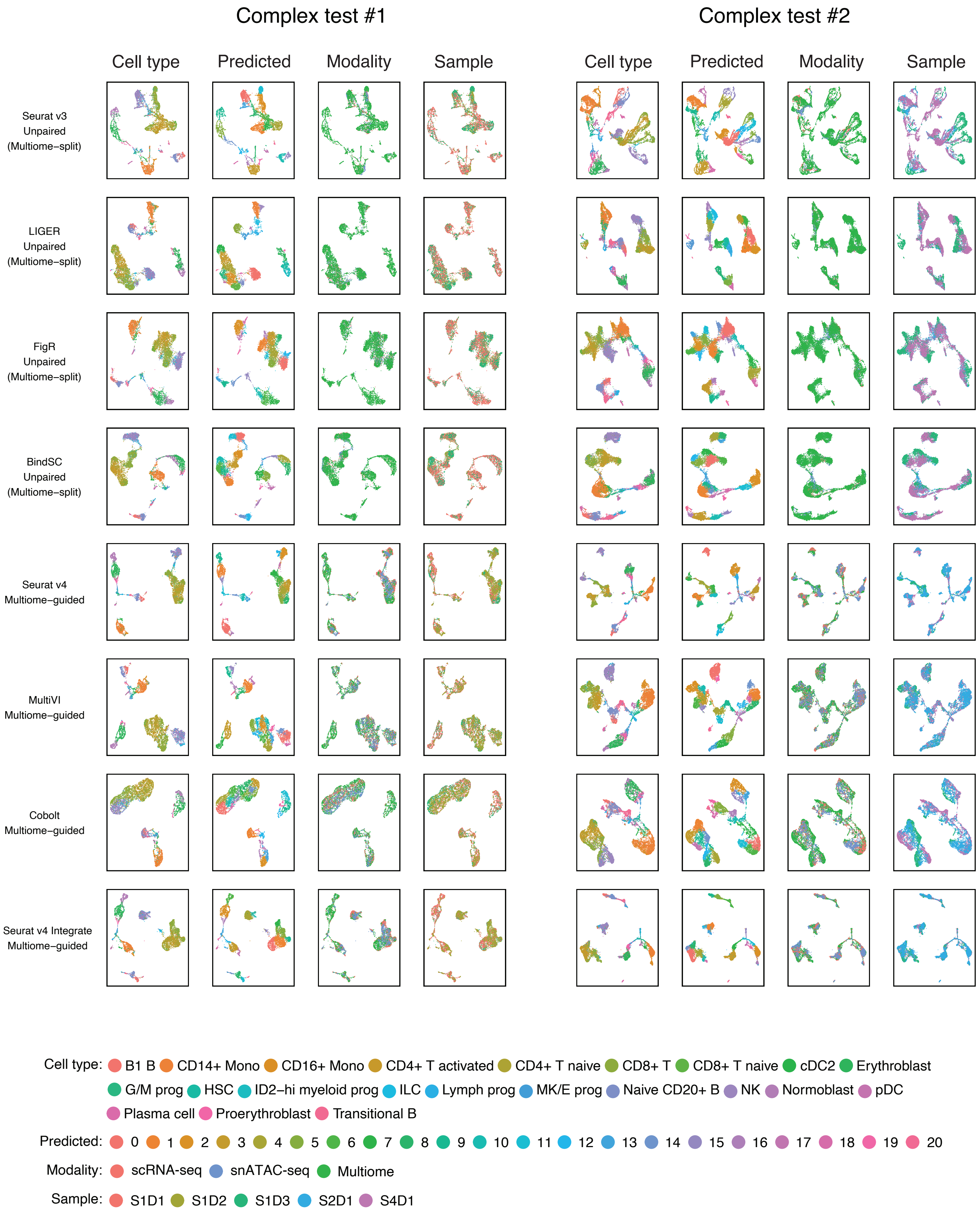


Supplementary Figure 14: UMAP plots for the BMMC-based simulations with complex batch effect challenge shown in Figure 4D.

#### Supplementary methods

We evaluated a total of seven methods. Each method was run according to the most relevant tutorial available. The source code is available on GitHub https://github.com/myylee/benchmark_sc_multiomic_integration.

**Multiome-guided integration methods**

Seurat v4 [1]

Each modality in the multiome dataset was processed with the standard approach. Specifically, RNA-seq profile was normalized with scTransform [2] and reduced to 50 dimensions using principal component analysis (PCA). ATAC-seq profile was processed with the latent semantic indexing (LSI) [3]. Specifically, the cell-by-peak matrix was normalized with term frequency-inverse document frequency (TF-IDF) and reduced to 50 dimensions using singular vector decomposition (SVD). The two separate latent embeddings were joined through the weighted-nearest neighbor (WNN) approach [4], using the top 50 principal components for the RNA-seq and the top 2-50 component for ATAC-seq, excluding the first component mainly correlated with sequencing depth. Single modality datasets were processed using the same standard workflow. Then, the raw data matrix was projected to the WNN-integrated space using supervised PCA [4] and supervised LSI. Normalized gene expression values were imputed using anchor-based approximation for the ATAC-seq cells. The multiome analysis was done https://satijalab.org/seurat/articles/weighted_nearest_neighbor_analysis.html#wnn-analysis-of-10x-multiome-rna-atac-1. ScRNA-seq was mapped following https://satijalab.org/seurat/articles/multimodal_reference_mapping.html#example-1-mapping-human-peripheral-blood-cells-1. We then adopted this framework to project the snATAC-seq dataset to the multiome reference.

Seurat v4 integrate

For the two simulations described in Figure 4D, the multiome dataset was composed of samples from multiple donors exhibiting batch effects. Before learning a joint representation for the cells using WNN, a within-modality integration was employed to mitigate batch effects. The RNA-seq profiles were integrated using canonical correlation analysis (CCA) described in https://satijalab.org/seurat/articles/sctransform_v2_vignette.html#perform-normalization-and-dimensionality-reduction-1. The ATAC-seq cells were integrated following steps described in https://stuartlab.org/signac/articles/integrate_atac.html. Then, the same WNN workflow and the supervised mapping of single-modality datasets were performed to project single-modality datasets to the batch-corrected space.

MultiVI [5]

MultiVI trains a variational autoencoder to learn a latent representation for cells in all three data types. Firstly, the two modalities of the paired datasets were horizontally stacked. Then, the paired and unpaired datasets were combined, with a modality column indicating which technology the cell belongs to. Features appearing in fewer than 1% of cells were filtered out. Then, the MultiVI model was trained with default parameters. Specifically, an autoencoder was trained for each modality using both the single-modality dataset and multi-modal dataset. Then, a symmetric Kullback-Leibler (KL) divergence loss was used to align the RNA-seq latent embedding and the ATAC-seq latent embedding of the paired cells. The model was trained using default parameters as described in https://docs.scvi-tools.org/en/stable/tutorials/notebooks/MultiVI_tutorial.html. The latent state of each cell was extracted using the converged model to represent the integrated cell state. The unmeasured RNA-seq profile of the unpaired ATAC-seq cells were inferred by passing the latent embedding through the learned decoder. When integrating datasets with additional batch labels, such as donor ID or site ID, these were included as covariates in the model.

Cobolt [6]

Cobolt also learns a latent representation of the cells using a variational autoencoder structure. Specifically, a modality-specific neural network was trained for each modality, using both unpaired and paired cells. For the paired cells, the latent representation of its

ATAC-seq profile and RNA-seq profile were multiplied to jointly define a cell's identity. A projection from the single modality latent embedding to the multi-modal latent representation was trained using the paired cells and later applied to the single-modality cells to ensure both unpaired and paired cells reside on the same space. Cobolt was run as described in the tutorial https://github.com/epurdom/cobolt/blob/master/docs/tutorial.ipynb.

#### Unpaired integration methods

##### Liger [7]

This method can integrate unpaired scRNA-seq and snATAC-seq datasets. The peak-count matrix was first converted into a gene activity matrix, aggregating reads mapped within 2-kb upstream or downstream of the transcription start site (TSS). Both ATAC-seq and RNA-seq profiles were normalized by total expression across each cell. Highly variable genes were selected using the second dataset, the RNA-seq profiles, using default parameters that select genes with variance greater than 0.1. Then the scaleNotCenter was used to scale the normalized data matrices. Joint matrix factorization was performed using optimizeALS with k = 20. Quantile normalization was performed on the resulting cell loading across RNA-seq and ATAC-seq profiles, allowing the datasets from two modalities to be integrated. Louvain clustering was then used with the goal to generate a specific number of clusters. UMAP loading was calculated using cosine distance between quantile normalized cell loadings, with 30 neighbors and a minimum distance of 0.3. We followed the tutorial at http://htmlpreview.github.io/?https://github.com/welch-lab/liger/blob/master/vignettes/Integrating_scRNA_and_scATAC_data.html.

##### Seurat v3 [8]

Seurat v3 integrates unpaired RNA and ATAC datasets using the canonical correlation analysis (CCA). RNA-seq was normalized by library size and log-transformed. Top 2,000 highly variable genes were identified and used for dimensional reduction with PCA. ScATAC-seq data was converted to gene activity matrix as described above and normalized in the same way as the RNA-seq data. Using the highly variable genes identified in the gene expression data as the features, anchors between RNA-seq and ATAC-seq were identified, and the two profiles were aligned using CCA. Canonical aligned components were used as integrated cellular loadings. Normalized gene expression was imputed by merging the expression of neighboring RNA cells. We followed steps from https://satijalab.org/seurat/articles/atacseq_integration_vignette.html.

FigR [9]

FigR projects scRNA-seq and snATAC-seq to one shared latent embedding using a similar workflow as Seurat v3. In addition, it computationally pairs cells from different modalities, creating a pseudo-multiome dataset for downstream analyses such as peak-gene pair identification. Specifically, the cell-peak count matrix was converted to gene activity matrix and normalized in the same way as the gene expression matrix. The top 5,000 most variable genes were identified using the gene expression matrix and the gene activity matrix individually, and the union of the two lists was obtained as the feature for downstream analysis. Canonical correlation analysis (CCA) was applied to align the ATAC and RNA profiles. L2 normalization was applied to the top 30 CCA components. OptMatch algorithms were used to pair the RNA and ATAC cells, using the 30 CCA component are latent embedding. Cells paired with multiple cells were removed, resulting in a 1-to-1 cell pairing. We implemented the preprocessing and CCA alignment of the unpaired datasets using functions in the Seurat package, and we used the OptMatch from the FigR GitHub page [9].

BindSC [10]

BindSC integrates unpaired single-cell datasets by estimating a cell-by-gene matrix Z for the snATAC-seq cells that maximizes the correlation between Z and the cell-by-peak matrix (Y) as well as between Z and the scRNA-seq data (X). The simultaneous similarity maximization is done in the latent embedding space through the bi-direction CCA. Firstly, 5,000 highly variable genes were selected using the RNA-seq profile. Then, the ATAC-seq profile was converted into the gene activity matrix as an initial approximation of the transformed matrix, Z. BiCCA was run with using default setting with lambda=0.5 and alpha=0.5, K=15, num.iteration=100, block.size=0. Through this process, Z was iteratively improved to better approximate the gene expression for the snATAC-seq cells while maintaining its similarity with the ATAC-seq profile. After training, a latent representation of the scRNA-seq and snATAC-seq cells was learned and could be used for clustering. Moreover, the imputation of gene expression was performed using impuZ and fold-change normalization is performed to scale the imputed RNA expression. Only the 5,000 highly variable genes used for integration were imputed. We followed the steps from https://htmlpreview.github.io/?https://github.com/KChen-lab/bindSC/blob/master/vignettes/mouse_retina/retina.html.
